# NOTCH-driven oscillations control cell fate decisions during intestinal homeostasis

**DOI:** 10.1101/2024.08.26.609553

**Authors:** Sonja D. C. Weterings, Hiromune Eto, Jan-Daniël de Leede, Amir Giladi, Mirjam E. Hoekstra, Wouter F. Beijk, Esther J. M. Liefting, Karen B. van den Anker, Jacco van Rheenen, Alexander van Oudenaarden, Katharina F. Sonnen

**Author notes:** These authors contributed equally (Sonja D. C. Weterings, Hiromune Eto).

## Abstract

Intestinal homeostasis requires tight regulation of stem cell maintenance and commitment to absorptive and secretory cells, two key intestinal lineages^1,2^. While major signalling pathways critical for this control have been identified^3^, how they achieve such a tight balance in cell type composition remains unclear. Here, we uncover dynamic expression of Hes1, a direct NOTCH target^3^, in intestinal stem and progenitor cells, and investigate its role *in vivo* and *in vitro*. A knock-in reporter^4^ reveals distinct, cell-specific period lengths in Hes1 oscillations that form a gradient along the crypt-villus axis. Whereas secretory precursors oscillate at low periods, absorptive precursors oscillate at higher periods before transitioning to a differentiated state. To test the function of different oscillation periods, we innovated a microfluidic system that modulates Hes1 oscillations in organoids. We find that varying the oscillation period modulates secretory cell differentiation: While 90-min oscillations promote Paneth cells, 130-min oscillations increase formation of other secretory subtypes. Moreover, low-period oscillations support stemness and a proliferative state. Our study provides the first clear evidence that information for tissue homeostasis in the intestine is encoded in the temporal dynamics of signalling components.

## Main

The intestinal epithelium renews every four to five days, maintaining stem and progenitor cells in the crypt and differentiated cells in the villus^5,6^. Dysregulation of intestinal renewal can result in intestinal atrophy, fibrosis and cancer^7^. Constant renewal and differentiation must therefore occur robustly across the lifespan of an organism. The secretory-to-absorptive cell ratio is crucial for efficient nutrient absorption and digestion, varying along the anteroposterior axis from approximately 1:6 in the small intestine to 1:3 in the large intestine^3,8–10^. Among secretory cells, goblet cells and Paneth cells are the most abundant, comprising 10–15% and 5% of the total epithelial cell population in the small intestine, respectively^11,12^. To maintain this cell-type composition, tissue-wide coordination of cell signalling is fundamental. Indeed, antagonistic gradients of WNT, EGF and NOTCH in the crypt and BMP in the villus determine the general organization of the crypt-villus axis^3^. Within the crypt, NOTCH signalling modulates critical cell fate decisions, including retention of stem cell fate and commitment to secretory or absorptive fate^1,2^. Over-activating NOTCH signalling constantly, such as through the overexpression of NICD (NOTCH intracellular domain), results in the loss of secretory cells and disrupts the vital balance between stem, absorptive and secretory cells^13^. The NOTCH target gene Hes1, which is highly expressed in the crypt, is crucial for these decisions as it represses several important differentiation factors^14,15^. In mice, Hes1 knockout is perinatally lethal due to brain defects; in the gut, it entirely skews the balance of secretory and absorptive cells towards the secretory lineage^14,15^.

Such static perturbations have not resolved critical questions about the maintenance of intestinal organisation, including how the intestine balances stem cell maintenance with differentiation, how the correct number of absorptive and secretory cells is specified, and what distinguishes secretory subtypes. Interestingly, NOTCH signalling acts dynamically in several tissues, rather than being “on or off”^16^. Complex dynamics can encode information, for instance, through signal duration or the periodicity of oscillations^17^. Members of the Hes family of transcriptional repressors, which are downstream of NOTCH signalling, have an intrinsic propensity to oscillate due to their own repressive feedback loop^18^. Hes1, for instance, oscillates during vertebrate segmentation and in the nervous system, muscle stem cells and pancreas^18–21^. While Hes1 dynamics have been proposed^22^ to occur in the intestine, and recently described for Hes6 in enteroendocrine precursors^23^, the nature and function of these oscillations remains untested *in vivo*. In addition, whether the mere presence or dynamic expression of such factors regulates cellular behaviour remains unclear.

### NOTCH-driven Hes1 oscillations occur in the small intestine

To investigate whether NOTCH-signalling dynamics occur in the small intestine, we used a knock-in mouse reporter line. The reporter is integrated into the Hes1 locus, a direct target of NOTCH signalling, to generate the fusion protein Hes1-Achilles (Fig. 1a)^4^. Homozygous Hes1-Achilles reporter mice were viable^4^ and showed no signs of intestinal insufficiency (i.e., weight loss or diarrhoea). We found similar cell type distributions within the crypt and villus in homozygous Hes1 reporter mice compared to wild-type mice (Extended Data Fig. 1)^4^, indicating that endogenous Hes1 tagging had no effect on intestinal patterning. As expected, Hes1-Achilles expression was restricted to the crypt region^24^ (Fig. 1b). It also showed a salt-pepper pattern in the crypt bottom where stem cells reside next to Hes1-negative Paneth cells. However, we observed more variability in Hes1-Achilles expression levels in cells located higher in the crypt. This observation suggested dynamic Hes1 expression in the small intestine.

**Fig. 1.**
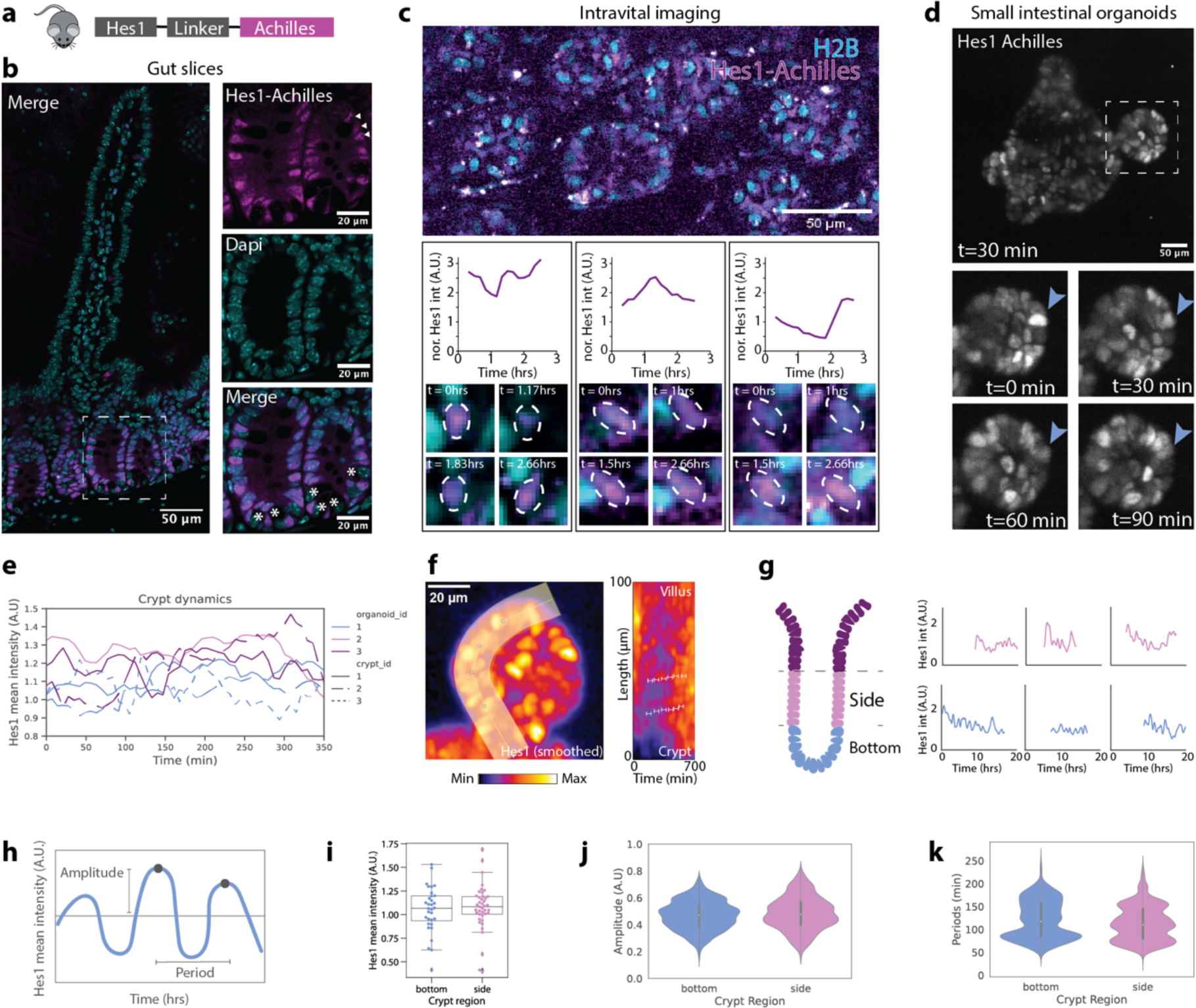
Hes1 is dynamic in small intestinal homeostasis. (a) Representation of the Hes1-Achilles reporter mouse line. (b) Representative image of Hes1-Achilles gut-slices counterstained with DAPI with magnification to show the crypt (right panels). Hes1-Achilles negative Paneth cells are indicated by asterisks. Arrowheads indicate varying Hes1-Achilles intensity at crypt sides. (c) Intravital imaging of the small intestine, representative image of Hes1-Achilles crossed with H2B-mCherry. Three example single-cell tracks (H2B normalized) are shown with a zoom-in of the associated imaging of these tracks over time. (d) Representative image of a Hes1-Achilles small intestinal organoid and (below) magnifications of crypt over time. Arrows highlight the same cell and its variable expression over time. (e) Hes1-Achilles intensity (mean-normalized) was measured in organoid crypts over time. (f) Kymographs from the crypt to the villus of cultured Hes1-Achilles small intestinal organoids were generated. Left panel: Zoom in of crypt with indicated kymograph line. Right panel: Kymograph over period of 10 hours and a length of 100µm Scalebar = 50µm (g) Left panel: Schematic representation of an organoid bud with bottom (blue) and side (pink) region highlighted. *Right panel:* Representative timeseries data of single-cell tracks in the three regions of the crypt, indicated with colour. (h) Schematic representation of an oscillation, amplitude and period indicated. (i) Quantification of Hes1-Achilles mean intensity. (j, k) Quantification of instantaneous amplitude (j) and period (k) by wavelet transform. Side: n=41 cells; bottom: n=32 cells. Tracks shorter than 7h were excluded.

To visualise Hes1 expression in real-time, we performed intravital imaging on the small intestine of mice expressing the Hes1-Achilles together with a fluorescent nuclear marker (H2B-mCherry). Phototoxicity and significant peristaltic movements of the gut restricted imaging quality and duration. Despite this, we tracked single cells in crypts over three hours and quantified Hes1-Achilles levels over time. We found that Hes1-Achilles was highly dynamic in these crypt cells (Fig. 1c, Supplemental Movie 1). Some cells exhibited rapid increases or decreases in Hes1-Achilles expression, while others displayed more gradual changes in Hes1-Achilles. We also observed cells at the crypt bottom that were constantly low in Hes1-Achilles expression, likely corresponding to Paneth cells (Extended Data Fig. 2). These observations suggest that Hes1 is dynamic in the small intestine *in vivo*.

To quantify these dynamics with higher spatiotemporal resolution, we generated intestinal organoids from the proximal small intestine of mice expressing the Hes1-Achilles together with a fluorescent nuclear marker (H2B-mCherry^25^ or iRFPnucmem^4^) (Fig. 1d). Intestinal organoids effectively mimic the cellular composition of the *in vivo* gut, including the differentiation into enterocytes and the two predominant secretory subtypes, Paneth and goblet cells^12^. Hes1-Achilles-expressing organoids showed a gradient of Hes1-Achilles expression, with the highest signal within the crypt, recapitulating the *in vivo* expression pattern^24,26^ (Fig. 1b, d).

To confirm a connection between the observed Hes1 dynamics and NOTCH activity, we first performed immunofluorescence (IF) staining against Hes1, which overlapped with reporter expression (Extended Data Fig. 3a). We then inhibited NOTCH signalling using DAPT (N-[N-(3, 5-difluorophenacetyl)-l-alanyl]-s-phenylglycinet-butyl ester), a small molecule gamma-secretase inhibitor. This treatment led to a drop in Hes1-Achilles levels within 30 min (Extended Data Fig. 3b, c), confirming that Hes1-Achilles expression is dependent on NOTCH signalling.

We live-imaged Hes1-Achilles organoids using an inverted light-sheet microscope, which allows high-resolution imaging for single-cell tracking with low phototoxicity. Hes1-Achilles was highly dynamic in most crypt cells by visual inspection (Fig. 1d, Supplemental Movie 2). Given that NOTCH signalling is known to cause spatial synchronization^27^, we examined crypt dynamics in addition to single-cell dynamics. When quantifying Hes1-Achilles dynamics in whole crypts, we observed varying expression levels over time (Fig. 1e). Kymographs generated along the crypt-villus axis indicated local, synchronized dynamics rather than the clear waves observed during somitogenesis^28^ (Fig. 1f). Subsequently, we quantified Hes1-Achilles expression in single cells, focusing on the crypt region. We divided cell positions into bottom and side regions to account for broad differences in cell-type composition (Fig. 1g). Single-cell tracks revealed that Hes1-Achilles expression displayed a variety of dynamics, oscillating in many cells in the bottom and side regions of the crypt (Fig. 1g and Extended Data Fig. 4). To quantify these oscillations, we used wavelet analysis, which fits wavelets of finite length centred around each time point along the track, providing an idea of “instant” oscillatory dynamics^29^. We determined the mean intensity, amplitude and the period of the detected oscillations (Fig. 1h). We did not observe differences in intensity or oscillation amplitude between bottom and side cells (Fig. 1i, j). Period quantifications revealed a broad range of periodicities between 75 and 200 min, with 3 main peaks at approximately 90, 130 and 170 min (Fig. 1k, Extended Data Fig. 4, 5). Thus, Hes1 expression oscillates with a range of periods in the crypts of intestinal organoids.

### Single-cell tracking in 2D organoids characterizes Hes1 oscillations

To fully capture the heterogeneity of Hes1 dynamics, we turned to a 2D organoid system (Fig. 2a). These simplified *in vitro* models are derived from intestinal stem cells or crypts and grow as a monolayer on a hydrogel-coated flat surface^30^. Due to its planar geometry, this system offers more straightforward automatic image analysis for single-cell tracking than the 3D organoid model, allowing higher-throughput quantification. The tissue self-organized into near-radially symmetric crypt-villus structures as seen in previous literature^31,32^. Verification with immunostaining confirmed that our system recapitulated differentiation into key cell types observed *in vivo* and in 3D organoid cultures, such as stem cells (Olfm4^+^), Paneth cells (Lyz1^+^), secretory cells (WGA^+^) and enterocytes (AldoB^+^) (Extended Data Fig. 6). Comparisons of these markers with Hes1-Achilles expression confirmed the absence of Hes1-Achilles in Paneth and secretory cells and high expression in stem cells. We observed graded Hes1-Achilles expression down the crypt-villus axis, correlating very well with the proliferation markers Sox9 and Ki67, extending to the TA (transit amplifying) zone^33^. Consistent with the literature^24^, Hes1-Achilles expression was absent in enterocytes (Fig. 2a, 1b).

**Fig. 2.**
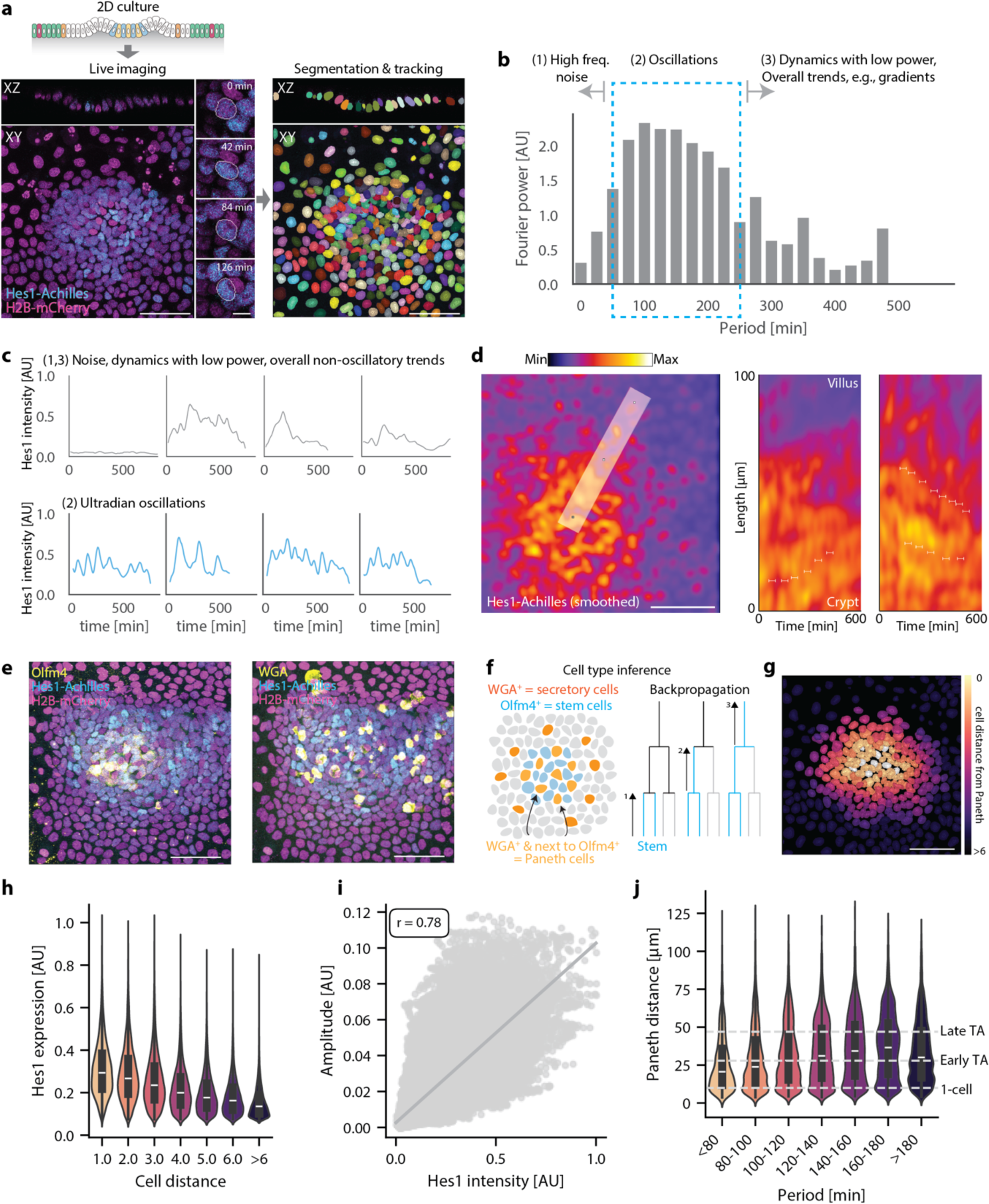
Single-cell tracking analysis of 2D intestinal organoids reveals an oscillation period gradient along the crypt-villus axis. Intestinal organoids expressing Hes1-Achilles and H2B-mCherry were cultured in 2D, followed by live-imaging and analysis. (a) Schematic representation, snapshots of live-imaging and segmentation result. Scale bars: 50 µm (overall), 5 µm (zoomed-in). (b) Mean Fourier spectrum of single-cell Hes1-Achilles intensity traces after removing fundamental and harmonic frequencies (full spectrum in Extended Data Fig. 7c). (c) Examples of Hes1-Achilles intensity traces over time, with raw data smoothed using a rolling mean over 3 timepoints. (d, left) Representative snapshot of a gaussian-smoothed 2D organoid expressing Hes1-Achilles, with a line along a crypt-villus axis from which kymograph was taken. Scale bar: 50 µm. (d, right) Two representative kymographs, brackets showing apparent synchrony (further examples in Extended Data Fig. 8a). (e) Immunofluorescence staining at the end of live imaging for Olfm4 (stem cell) and WGA (secretory cell). Scale bar: 50 µm. (f) Schematic representation of cell type inference to define secretory, stem and Paneth cells. (g) Segmented nuclei representing distance from Paneth cells, scale bar 50 µm. (h) Violin plot of Hes1-Achilles expression vs cell distance from Paneth cells. (i) Scatter plot of amplitude vs Hes1-Achilles intensity from wavelet analysis, with Pearson correlation coefficient, r, calculated by linear least-squares regression, p = 0.0. (j) Violin plot of distance from Paneth cells vs period from wavelet analysis. For panels (h, j) n=1901 tracks, N=3 biological replicates, n_inst_ = 147959 instantaneous measurements. Normal distribution was assessed by Shapiro Wilk test, p<0.05 for all datasets. Significance was tested by Mann-Whitney U test, with all conditions statistically different from each other, p<0.05.

We conducted 24-hour live-imaging at high spatial resolution to capture Hes1-Achilles dynamics in individual cells (Fig. 2a, Supplemental Movie 3). For high-throughput analysis, we developed an automatic nuclear segmentation and tracking pipeline to quantify Hes1-Achilles signals in individual cells (Fig. 2a, Supplemental Movies 4 and 5, see Methods for details). Given that Hes1 displays 1.5 – 3 h period oscillations in other systems^34^, we excluded tracks shorter than 8 hours. This approach therefore quantifies *ultradian* oscillations, i.e., with periodicity occurring from one to several hours, with sufficient confidence. This filtering resulted in a complete track length distribution of 8–20 hours (Extended Data Fig. 7a, b).

The analysis of single-cell tracks revealed a diverse range of Hes1-Achilles dynamics, characterized by varying intensity levels over time. We applied Fourier transformation to the intensity traces to identify frequency peaks characteristic of oscillatory systems. After removing contributions resulting from fundamental and harmonic frequencies related to track lengths (see Methods), the mean Fourier spectrum across all tracks unveiled a broad but distinct peak in the 75-250 min range (Fig. 2b). In addition to short intensity fluctuations and long-term increases or decreases in Hes1-Achilles levels, most cells exhibited oscillatory behaviour with distinct periodicity (Fig. 2b,c). These oscillations show similar periodicity to other systems where dynamic expression plays a critical coordinating role^34^. These data reveal that Hes1 expression shows ultradian oscillatory behaviour in the small intestine.

### Synchronisation of Hes1 oscillations

To analyse intercellular synchronization in more detail, we generated kymographs along lines from crypt centres towards villi, which revealed various neighbouring vertical lines of Hes1-Achilles expression along space, supporting the presence of tissue-wide coordination (Fig. 2d, Extended Data Fig. 8a). To quantify local synchrony of Hes1-Achilles oscillations between neighbouring cells, we calculated the Kuramoto order parameter between a cell and its direct neighbours (Extended Data Fig. 8b, c). The median order parameter across the tissue was 0.39 for all tracks, which indicates the oscillations of neighbouring cells are not always synchronized, explaining why such periodicity has been difficult to observe *in vivo*. However, we occasionally observed spikes in the instantaneous order parameter, reaching values near 1. These transient increases in synchrony also appeared as sporadic patches in the live imaging data (Extended Data Fig. 8c, Supplemental Movie 6).

Interestingly, we noticed Hes1-Achilles expression peaks in daughter cells after division (Supplemental Movie 7, Extended Data Fig. 8d, e). We observed a significant dip in Hes1-Achilles expression at mitosis, likely due to nuclear envelope breakdown and the subsequent release of Hes1-Achilles into the cytosol. Examining the phase of Hes1-Achilles oscillations around cell division, we found that phases synchronised approximately 1.5 to 1 hour before mitosis and were maintained until about 1 to 1.5 hours after division (Extended Data Fig. 8f, g). These findings suggest correlation between cell division and the synchronous Hes1 behaviour of neighbouring cells.

### Backtracking reveals a period gradient along the crypt-villus axis

To elucidate the spatial distribution of oscillatory Hes1 behaviours in the intestine, we developed an analysis framework to define the crypt centre in our live imaging data. By backtracking from immunofluorescence staining of cell type markers, we defined stem, Paneth and secretory cells (Fig. 2e,f, see Methods). We then calculated the distance from the crypt centre as the number of cells away from Paneth cells (Fig. 2g, Extended Data Fig. 9a).

Quantifying Hes1-Achilles dynamics within this framework, we observed that the gradient of Hes1-Achilles intensity decreased with distance from Paneth cells^24^ (Fig. 2h), supporting the validity of our approach. Wavelet analysis revealed an amplitude gradient correlating well with the intensity gradient (Fig. 2i, Extended Data Fig. 9b). The wavelet power spectra for individual tracks showed varied oscillatory patterns: some tracks oscillated at a single period (Extended Data Fig. 10a), while others exhibited changes in period or had multiple, nested periods (Extended Data Fig. 10b, c), indicating complex oscillatory behaviour.

The distribution of periods changed significantly with increasing cell distance from Paneth cells (Extended Data Fig. 9c). Notably, lower periods existed almost exclusively near Paneth cells (Fig. 2j). Periods less than 80 min and those between 80-100 min peaked at around 12 µm (approximately one cell distance from Paneth cells) and had a median around 22-25 µm (within two cells). This conclusion agrees with periodicity quantified in the crypts of 3D organoids (Fig. 1k): Cells near Paneth cells had a prominent peak at 90 min but showed a broader distribution extending further up the crypt-villus axis. Thus, a period gradient was detectable, with periods lengthening as the distance from Paneth cells increased.

### Cell type inference reveals cell type-specific Hes1 dynamics

To determine which cells exhibited specific oscillatory behaviour, we defined cell types based on IF staining after live imaging. In addition to stem, secretory and Paneth cells (Fig. 2e, f), we classified immature progenitor cells (IMPCs, precursors to secretory cells), enterocytes and TA cells (Fig. 3a, b, further details in methods). To distinguish the TA cell states more clearly, i.e., more progenitor-like or more differentiated, we subdivided them into Early (up to three cells) and Late TA cells (from four cells onwards) based on their positions relative to Paneth cells (Fig. 3a, b, further details in methods). Based on this cell type classification, we reanalysed our tracks. Different cell types were positionally distributed, as expected, according to their distance from Paneth cells: stem cells were mostly at the 1-cell position, and enterocytes were further away (>6 cells) (Extended Data Fig. 11a). Hes1-Achilles expression was highest in stem cells and followed a gradient that decreased from Early to Late TA cells and then to enterocytes, while secretory cells showed low Hes1-Achilles expression (Fig. 3c, Extended Data Fig. 11b-e).

**Fig. 3.**
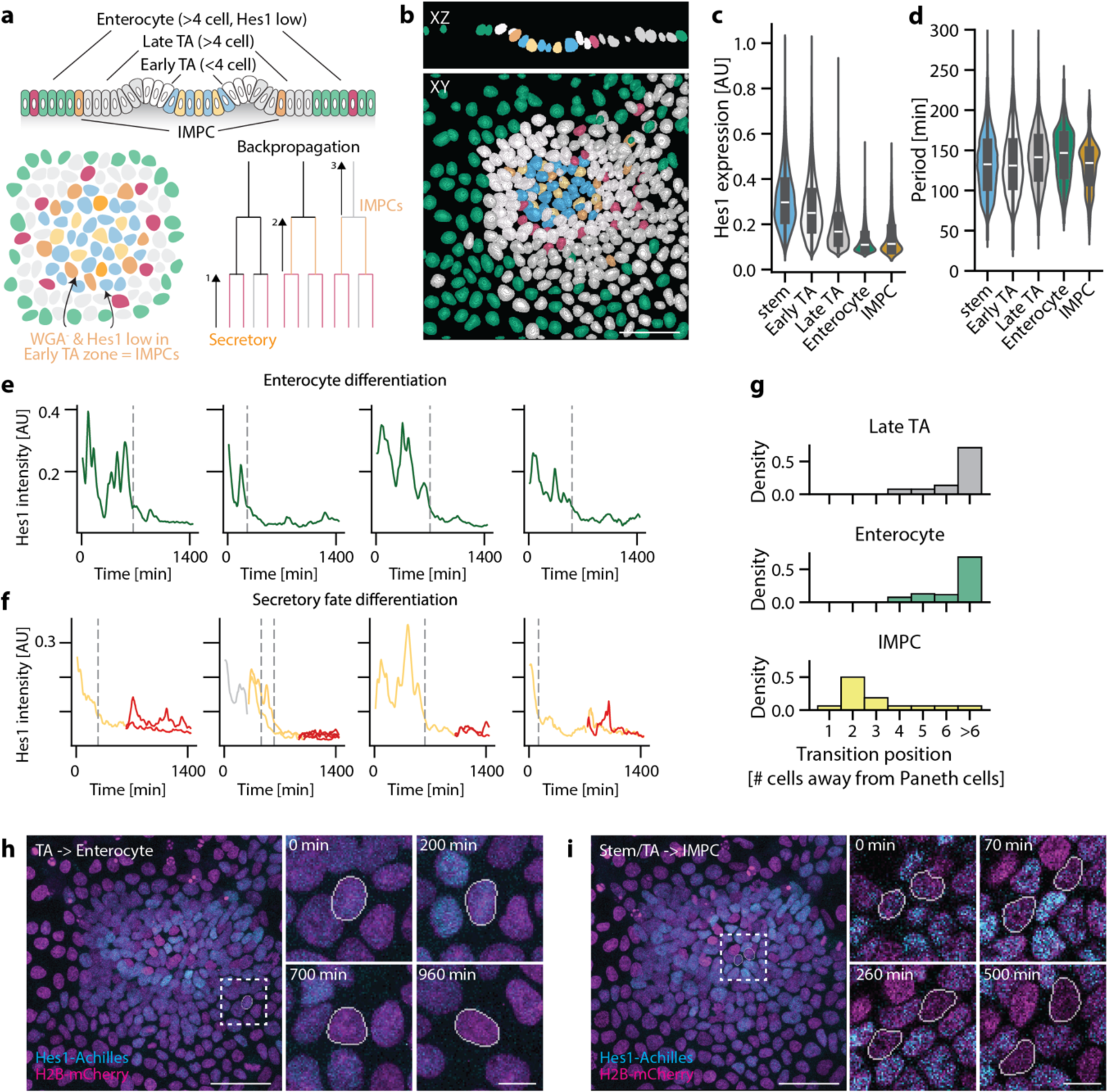
Different intestinal cell types display distinct Hes1 dynamics. Intestinal cell types were defined for all single-cell tracks from 2D organoid live imaging, followed by analysis. (a) Schematic representation of cell-type inference and (b) segmentation showing resultant classification. Scale bar: 50 µm. (c) Violin plot of Hes1-Achilles expression vs cell type. (d) Violin plot of Hes1-Achilles expression vs period from wavelet analysis. For panels (c, d), n=1901 tracks, N=3 biological replicates, n_inst_ = 147959 instantaneous measurements. Normal distribution was assessed by Shapiro Wilk test, p<0.05 for all datasets. Significance was tested by Mann-Whitney U test, with all other cell types compared against stem cells. All comparisons were statistically significant, p<0.05. Representative Hes1-Achilles intensity traces for (e) enterocyte differentiation and (f) secretory fate differentiation, grey dotted lines indicating moment of transition (see Methods). (g) Histogram of Hes1-Achilles high-to-low transition position per cell type, n_enterocyte_ = 190, n_Late TA_ = 243, n_IMPC_ = 16. Representative images and snapshots of (h) enterocyte differentiation, and (i) secretory cell differentiation in 2D organoids expressing Hes1-Achilles and H2B-mCherry. Scale bars: 50 µm (overall), 5 µm (zoomed-in).

Next, we re-evaluated our wavelet analysis and detected significant differences in oscillation period between cell types (Fig. 3d). Stem cells and Early TA cells had lower medians (132 min for both) compared to Late TA (142min) and enterocytes (149min). The latter included cells transitioning from TA cells to enterocytes during live imaging and showed a prominent peak at 170 min period. This is consistent with the period gradient observed earlier. Amplitudes also correlated well with intensity (Extended Data Fig. 11f). IMPCs had a median at 133 min, similar to stem or Early TA cells, and also a distinct peak at around 90 min (Fig. 3d).

To investigate cells transitioning to the absorptive or secretory lineage in detail, we focused on cells showing a Hes1-Achilles expression arrest midway through the tracks (Fig. 3e-i). Most progenitor-to-enterocyte transitions occurred far down the crypt-villus “differentiation” gradient (Fig. 3e, g, h, Supplemental Movie 8). This gradient indicates that cells gradually decreased in Hes1-Achilles intensity and transitioned to a low-Hes1-Achilles, differentiated state as they moved away from the crypt centre. In the second case, most IMPC transitions occurred very early in the crypt-villus axis^35^, peaking at the 2-cell position, with some even occurring directly adjacent to Paneth cells (Fig. 3f, g, ii, Supplemental Movie 9). Most cells in this vicinity were Hes1-Achilles-high cells (Fig. 2h, Extended Data Fig. 11a-e). These cells therefore transitioned early from a high-expression oscillatory state to a Hes1-Achilles-low state, committing to the secretory lineage.

When analysing the cell-type dependence of local synchronization, we found that Kuramoto order parameters between cells of each cell type and their neighbours were comparable with a slight trend to higher values for transitioning enterocytes (Extended Data Fig. 11g). Finally, we quantified cell cycle length for each proliferating cell type (Extended Data Fig. 11h). Stem and TA cells cycled on average approximately every 11 h^35^, while IMPCs had long cycling times of 18.3 h. When aligning tracks to the point of cell division, all cell types showed a dip in Hes1-Achilles expression at division followed by a peak (Extended Data Fig. 11i). Even though IMPCs were mostly Hes1-Achilles low throughout their lifetime (Extended Data Fig. 11e), they still divided once more after a drop in Hes1-Achilles signal to give rise to secretory cells.

Thus, our data suggest that stem and progenitor cells show lower-period Hes1 oscillations. Moreover, cells commit to a secretory fate in the low-period region of the Hes1 oscillation gradient, where IMPCs oscillate with low and mid-range periods before transitioning. Conversely, enterocyte precursors oscillate at higher periods before transitioning in the high-period region of the Hes1 gradient.

### Modulation of NOTCH signalling dynamics

Next, we sought to modulate dynamics, rather than absolute expression levels, to determine the functional effects of different oscillation periods. To this end, we established a microfluidic system for the on-chip culture of 3D organoids (Fig. 4a, Extended Data Fig. 12a-c). This system allows modulation of culture conditions over time whilst simultaneously monitoring Hes1-Achilles dynamics by fluorescence microscopy. Marker expression for stem cells (LGR5-DTR-GFP^36^), enterocytes (CK20) and apoptosis (cleaved Caspase-3) verified proper organoid differentiation and survival in the microfluidic system (Extended Data Fig. 12d,e).

**Fig. 4.**
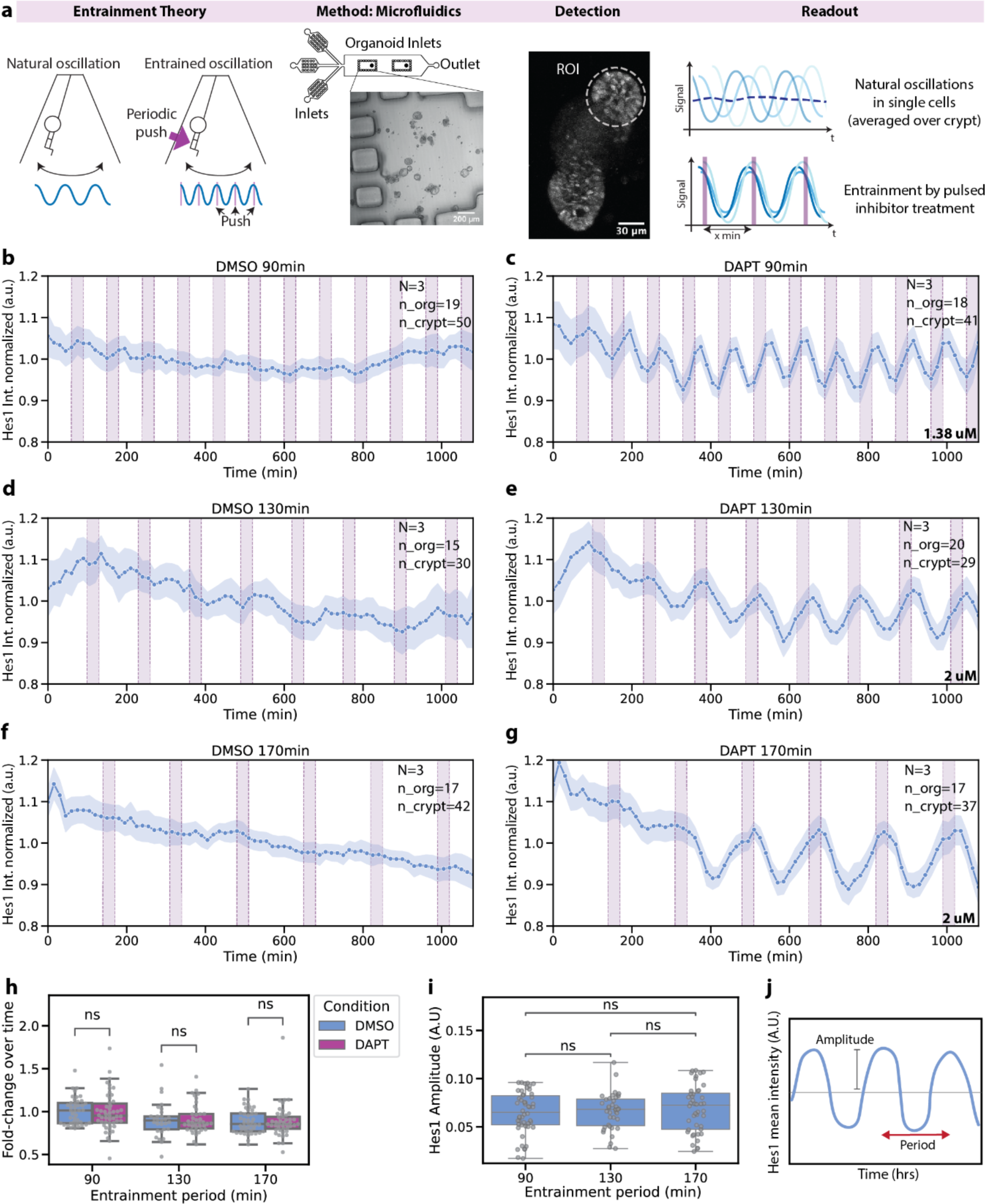
Entrainment using microfluidics modulates signalling dynamics in organoids. (a) Schematic overview including the entrainment theory, the microfluidic chip, including a representation of organoids cultured within the chip, crypt dynamic detection of Hes1-Achilles intensity overtime and our hypothesis of the experimental read-out Organoids were entrained to a 90-130- or 170-min period for 20-24 hours. (b-g) Entrainment of Hes1-Achilles oscillations to periodic pulses, 90-(b, c) 130-(d, e) and 170-min (f, g) of the NOTCH inhibitor DAPT (concentration indicated). Quantification of (normalized) Hes1-Achilles signal in organoid crypts. Experiments (N = independent experiments, n_org = individual organoids, n_crypt = individual crypts) were combined using the external force (pink bars) as an objective time reference. (h) Quantification of Hes1-Achilles absolute intensity overtime by calculating the fold-change over time, comparing the intensity of the first 20 timepoints to the last 20 timepoints of each condition. Comparison between control (DMSO) and entrainment (DAPT) Hes1-Achilles fold change overtime. Independent t-test was performed per condition. (i) Quantification of amplitude by wavelet transform and comparison between entrainment periods. Mann-Whitney U was performed, with Bonferroni multiple sample correction. (j) Schematic representation of a Hes1 oscillation highlighting the period, which was the only feature changed in experiments shown this Figure.

To control oscillations, we used an entrainment approach, which synchronises endogenous oscillations with periodic external pulses (Fig. 4a)^36–38^. We achieved synchronisation of Hes1-Achilles oscillations through periodic application of DAPT^38,39^. We aimed at modulating oscillations to 90, 130 and 170 min, spanning the range of the Hes1-Achilles period gradient we had detected earlier. We selected a DAPT concentration that provided only partial inhibition of Hes1-Achilles when applied continuously to subtly modulated signalling activity (Extended Data Fig. 12f). Furthermore, we initially adjusted DAPT concentrations according to each period — 90 min, 130 min, and 170 min — to ensure the cumulative dose remained consistent across all conditions. To monitor drug pulses, we tracked the flow of DMSO or DAPT into the microfluidic chamber using the fluorescent dye cascade blue (Extended Data Fig. 12g, h). We then analysed Hes1-Achilles dynamics by quantifying reporter expression within crypts (Fig. 4a).

Periodic DAPT pulses synchronised endogenous Hes1-Achilles oscillations with the external DAPT rhythm across all entrainment periods (Fig. 4b-g, Supplemental Movie 10). We quantified the amplitude and fold-change of absolute Hes1-Achilles levels to ensure there were only changes in Hes1-Achilles-oscillation period. While 90- and 130-min pulses did not alter amplitude or absolute Hes1-Achilles levels, significant changes in amplitude were observed with 170-min pulses (Extended Data Fig. 12i-k). Therefore, we lowered the DAPT concentration to promote 170-min entrainment conditions, resulting in entrainment without altering amplitude or absolute Hes1-Achilles levels (Fig. 4h, i). Thus, our microfluidic system effectively modulates the period of NOTCH-driven oscillations in intestinal organoids (Fig. 4j).

### Bulk RNAseq indicates changes in cell type composition upon modulation of oscillations

To identify potential subtle changes in cell type composition upon modulation of Notch-driven oscillations, we performed bulk RNA sequencing (bulk RNAseq) after entraining oscillations to 3 different periods (90-, 130- and 170-min) for 3 different durations (24-, 48- and 72-hours) (Fig. 5a). Organoids were harvested at the same ‘developmental age’ (i.e., 4 days after passaging) to reduce potential variations in cell type composition based on organoid age (Extended Data Fig. 13a, b). We compared results based on the duration of on-chip culture to remove any bias, i.e., we for instance compared all 24 h DAPT samples to a 24 h DMSO control.

**Fig. 5.**
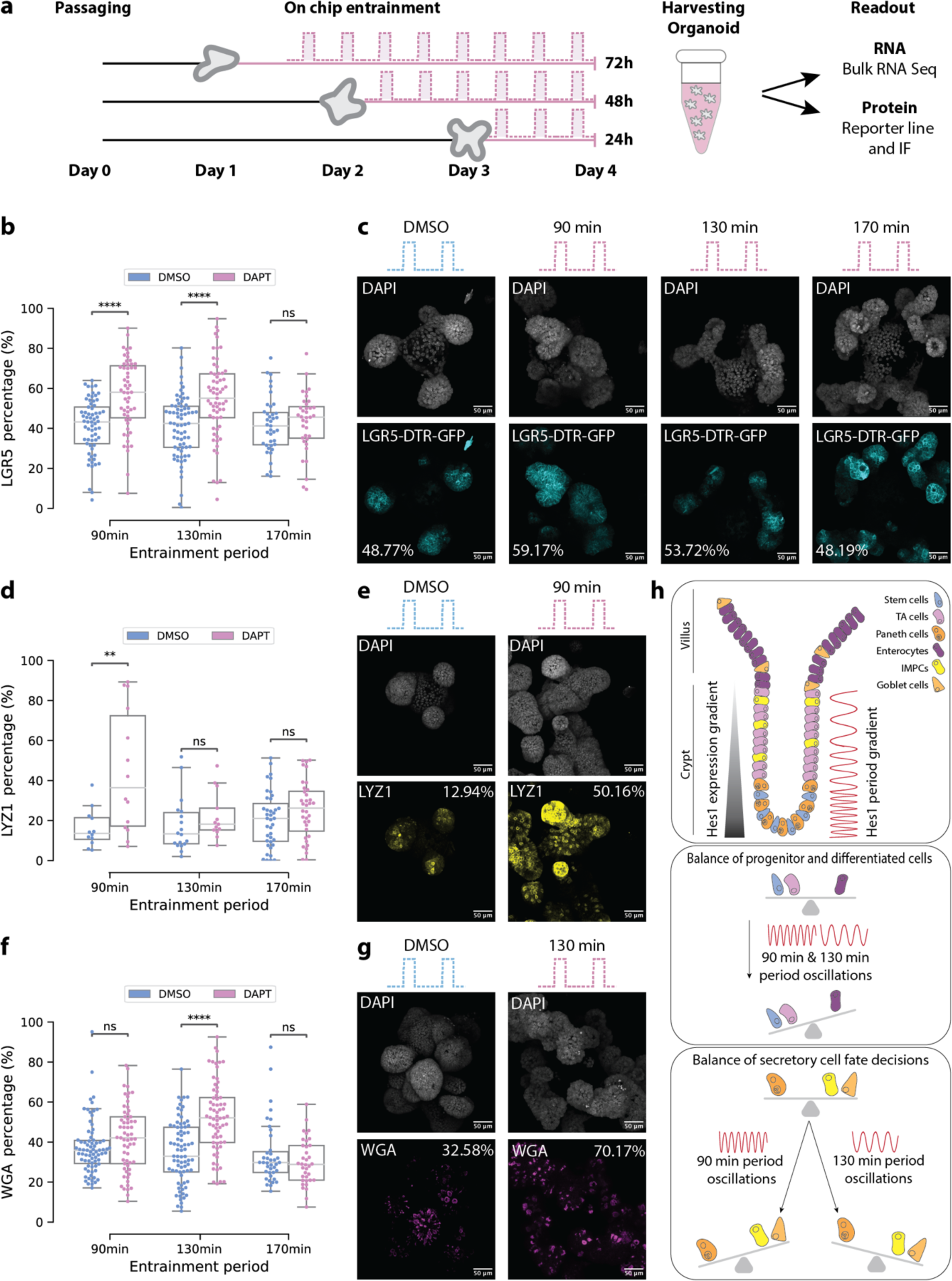
NOTCH signalling dynamics drive crypt maintenance. (a) Schematic representation of experimental design. Organoids were cultured for one, two or three days in a normal culture dish and entrained within the microfluidic chip for 72, 48 or 24 hours respectively, for all 3 entrainment periods: 90-, 130-,170-min. All organoids were harvested at day 4 and send for RNA bulk sequencing or protein analysis. (b-g) Quantitative analysis of cell percentage per organoid compared to control (DMSO) for 90-,130- and 170-min entrainment for 72 h and representative images including percentage. Cell percentages tested for LGR5 (b, c), LYZ1 (d, e), WGA (f, g). N= minimum of 3, and n_org = minimum of 14 organoids. Normal distribution was calculated using the Shapiro Wilk test and significance was tested by independent t-test or Mann-Whitney U test depending on normal distribution. ns: 5.00e-02 < p <= 1.00e+00, *: 1.00e-02 < p <= 5.00e-02, **: 1.00e-03 < p <= 1.00e-02, ***: 1.00e-04 < p <= 1.00e-03, ****: p <= 1.00e-04. (h) Model of Hes1 oscillations determining cell proliferation and differentiation in the small intestine.

When investigating expression of known cell type markers, we indeed found subtle effects depending on entrainment conditions (Extended Data Fig. 14). Both 130- and 170-min entrainment generally showed similar expression changes. There was an increase in Paneth cell markers after 72 h of 90-min entrainment (90min(72h)-entrainment) as well as a slight decrease in enterocyte markers after 130min(72h)-entrainment. In the latter condition, we also found an increase in markers for fetal-like stem cells. Fetal-like stem cells (markers *Msln, Ly6a, Clu, Anax1*) are found in the gut during embryonic development^40^, regeneration of adult intestines^41^, in organoids after passaging^42^ and occasionally in the *in vivo* small intestine (Extended Data Fig. 13).

To more accurately extract information on cell type composition, we applied the Latent Dirichlet Allocation (LDA) approach to our bulk RNAseq data, using the single-cell RNAseq (scRNAseq) data presented in Bues et al. (2022)^40^ (Extended Data Fig. 15). Our approach performs deconvolution of bulk RNAseq data into distinct cellular states^40^. Cell-type changes between entrainment periods and controls (Extended Data Fig. 15e-n) revealed an over-representation of Paneth cells after 90min(72h)-entrainment. For 130min(72h)-entrainment, we observed a decrease in enterocytes and an increase in progenitor cells, including fetal-like stem cells. In addition, there were indications of a decrease in enteroendocrine cells after 170min(72h)-entrainment, even though expression of enteroendocrine markers was low, potentially leading to large variations in observed results (Extended Data Fig. 15e-g, j). Thus, we found indications for changes in stem cell maintenance and differentiation depending on oscillation period.

### NOTCH-driven oscillations modulate crypt maintenance

To confirm whether oscillation period regulates cell type composition, we quantified cell types by staining for cell type markers using IF upon entrainment. We focused on conditions with the most potent effects, as indicated by bulk RNAseq data (24 h: 90 and 130 min; 72 h: 90, 130 and 170 min) (Fig. 5a). WT or Lgr5-DTR-GFP organoids^43^ were entrained, then stained for WGA (secretory cells), LYZ1 (Paneth cells), SOX9 (progenitor zone) and SCA-1 (fetal-like stem cells). To quantify changes in cell type abundance after entrainment from a large number of images, we performed superpixel analysis (Extended Data Fig. 16a). To ensure the accuracy of our study on homeostatic organoids, we excluded small, non-budded organoids from our analysis.

As a control for our microfluidics experiments, we continuously treated organoids with the average DAPT concentration equivalent to the cumulative dose of microfluidics experiments (0.48 µM for 90- and 130min entrainment, while in 170-min entrainment only 0.35 µM was used). This treatment resulted in no changes in morphology or cell type distribution (Extended Data Fig. 17), supporting the conclusion that any observed effects upon entrainment were due to changes in oscillation periods rather than absolute NOTCH signalling levels.

Whereas entrainment for 24 h showed no apparent changes in cell type composition, presumably due to the required time for differentiation and marker expression (Extended Data Fig. 16b-g), we determined two main effects after 72 h of entrainment.

First, 90min(72h)- and 130min(72h)-entrainment increased the relative amount of LGR5^+^ stem and SOX9+ progenitor cells in organoids, while 170min(72h)-entrainment did not (Fig. 6b,c, Extended Data Fig. 16h). Corresponding to a shift towards undifferentiated crypt cells, organoids contained fewer enterocytes (determined as LGR5-, SOX9- and WGA-superpixels) (Extended Data Fig. 16i, j). In contrast to the bulk RNAseq data, we could not confirm a significant effect on fetal-like stem cells, even though we found a trend towards higher percentages of SCA-1^+^ cells under all entrainment conditions (Extended Data Fig. 16k). Thus, our data suggest that low (90- to 130-min) period oscillations shift the organoid cell balance towards undifferentiated stem and progenitor cells.

Second, we identified a period-specific effect on the number of secretory cells. Consistent with the bulk RNAseq data, we found that 90min(72h)-entrainment increased the percentage of LYZ1^+^ Paneth cells (Fig. 5d, e). In contrast, no changes in the percentage of Paneth cells were detected in other entrainment periods (Fig. 5d). When analysing the broader secretory lineage marker WGA, we found an increase in WGA^+^ cells specifically in the 130min(72h)-entrainment. 90min(72h)-entrainment showed a trend towards higher percentages (p = 0.052), which is consistent with the increased percentage in Paneth cells as one of the secretory subtypes (Fig. 5f,g, see above). Conversely, 170min(72h)-entrainment did not affect this percentage. This suggests that 90-min oscillations promote Paneth cell differentiation, while 130-min oscillations promote differentiation of secretory cells other than Paneth cells.

## Discussion

Here, we find that Hes1, a direct target of NOTCH signalling, oscillates in the intestine with the periodicity of 90 min at the crypt bottom and 170 min towards the villi. Modulation of the oscillation period using microfluidics-based perturbation resulted in significant changes in cell type composition (Fig. 5h). While our microfluidic approach targets entire 3D organoids, which leads to an increase in synchronization of oscillations between neighbouring cells for all entrainment conditions, we identified period-specific effects. This highlights that (1) it is not intercellular synchronization that modulates cell type composition and (2) specific periods promote specific changes in cell identity.

While enterocyte precursors slow down in Hes1 oscillations towards high periods, secretory precursors show low- and mid-range-period oscillations before transitioning. We show that entraining oscillations to 90-min periods promotes Paneth cell formation, while 130-min oscillations support the appearance of secretory cells other than Paneth cells, presumably goblet cells. The period distribution of secretory precursors with few cells oscillating at 90 min and a majority at 130 min is consistent with differences in Paneth (5 %) and goblet cell (10 – 15 %) numbers in the small intestine. This indicates that dynamics play a role in regulating intestinal cell type composition, in particular by modulating secretory cell differentiation. In enteroendocrine progenitors, another secretory subtype, oscillations in the Hes6-target gene Ascl1 have recently been observed, indicating potential oscillations of other Hes genes in the intestine^23^. Since we used a general NOTCH pathway inhibitor to entrain oscillations, our microfluidic system modulates all NOTCH-dependent target genes. Our approach therefore provides unprecedented insight into the functional effects of periodic NOTCH-pathway activity in the intestine. How individual Hes genes contribute to differentiation and interact with Hes1, and its target genes (Atoh1, Ascl1) are interesting directions for future research.

Enterocyte precursors have been shown to proliferate more often than secretory precursors before terminally differentiating, which is thought to regulate intestinal enterocyte abundance^35^. We find that enterocyte precursors continue oscillating in Hes1 throughout the crypt, with increasing oscillation periods towards the crypt top. Interestingly, entraining oscillations to periods of 90 and 130 min, but not 170 min, promotes stemness and a proliferative state. This indicates that low-period oscillations prevent enterocyte differentiation in the bottom of the crypt. This agrees with similar findings in other contexts, such as cells in brain and breast tissue, where Hes oscillations correlate with proliferation^44–46^. Our data suggest that it is not the presence of oscillatory factors *per se,* but specific oscillation periods that regulate the number of proliferative cells in the gut. Moreover, we show that Hes1 oscillations are correlated with cell cycle dynamics, with cells consistently showing a peak of Hes1 expression after cell division. Future studies should reveal whether NOTCH-driven oscillations are functionally linked with cell proliferation^47^ or indirectly modulate a proliferative state by promoting stemness.

Our findings are consistent with a model in which oscillations in the crypt modulate intestinal patterning (Fig. 5h). In this framework, a gradient of periodicity along the crypt-villus axis maintains stemness and a proliferative state towards the bottom of the crypt. This way, enterocyte precursors are maintained in an undifferentiated state throughout the crypt length to expand in cell numbers. Moreover, stem cells differentiate towards secretory cells at the crypt bottom with a 90-min oscillation period promoting Paneth cell formation and 130-min promoting goblet cell formation.

These results will drive further investigations into how oscillations are regulated at the cellular level and how oscillations interact with other signalling pathways, including morphogen gradients^48^ and intestinal ERK dynamics^49,50^, to modulate intestinal cell type composition along the anteroposterior axis of the intestine^10^. This also has direct implications for understanding homeostasis of other tissue types and non-physiological conditions such as inflammation, regeneration and cancer.

## Methods Summary

### Extended Methods

#### Ethics

This study complies with all relevant ethical regulations. All animal experiments were approved by the Animal Experimental Committees at the Hubrecht Institute and the Netherlands Cancer Institute, and the Dutch Central Authority for Scientific Procedures on Animals (CCD). All animals were bred and housed under standard conditions (humidity 50–60%, 22–23 °C, inverted 12/12 h light/dark cycle, water and standard chow ad libitum) at the Hubrecht animal facility or at the Netherlands Cancer Institute facility in Amsterdam, the Netherlands.

### Mouse model

Hes1-Linker-Achilles or Hes1-Achilles (Hes1^tm1(Link–Achilles)Kfs)4^ crossed with H2B-mCherry (R26-H2B-mCherry, CDB0204K)^25^ or iRFPnucmem (Gt(ROSA)26Sor^tm1(iRFPnucmem)Aau^)^4^ were used for this project. C57BL/6 mice were used as controls. The H2B-mCherry mouse line was kindly provided by the Riken Center for Biosystems Research, the iRFPnucmem mouse line by Alexander Aulehla (EMBL Heidelberg) and the Lgr5tm2(DTR/EGFP)Fjs mouse line by Fred de Sauvage (Genentech)^43^. Mice were genotyped using standard PCR procedures, for primers see Supplementary Table 1.

### Paraffin sectioning of gut rolls and immunofluorescence

Two males and two virgin females of 8 weeks from both the Hes1-Achilles line and C57BL/6 control were used. Small intestine was isolated (from beneath the stomach to the caecum) and immediately flushed with formalin 4%. The tissue was rolled and stored in formalin 4% for 1 day at RT. This was followed by dehydration using a series of ethanol solutions (30%, 50% and 70%). Next, the gut-rolls were soaked in Xylene, embedded in paraffin and made into 5 um coupes. For staining the gut roll microscope slides, the slides were soaked in Xylene, 100% ethanol, 95% ethanol, 70% ethanol, 50% ethanol and rinsed with running cold tab water. For antigen retrieval the slides were cooked in a citrate buffer (sodium citrate (8.2mM) and citric acid (1.8mM); pH=6). 5% Goat Serum (cat. no. ab7481) in 0.3%-Triton PBS was used for blocking and permeabilization. Primary (overnight (ON) at 4°C) and secondary antibodies (3h at room temperature (RT)) were used in 0.1%BSA PBS and are listed in Supplementary Table 2. After staining, gut roles were mounted with Vectashield (cat. no. H1200NB). The samples were imaged using a SP8 confocal microscope (Leica Microsystems, 40x oil objective, numerical aperture (N.A.) = 1.3).

### Vibratome sections of small intestine

One virgin female of the Hes1-Achilles background was used for this previously described method^51^. In short, the small intestine was isolated (from beneath the stomach to the caecum) and immediately flushed with formalin 4%. The tissue was stored in formalin 4% protected from light (to maintain endogenous fluorescent signal) for 1 day at RT. 1cm of tissue was embedded in 4% of low melting agarose (cat. no. 16520100) and sections of +/- 150 µm were made using a microtome. Microtome settings included: velocity of 80mm s-1, knife amplitude of 0.70mm and vibration frequency: 65Hz. Tissue slices were blocked and permeabilized with 5% Goat Serum in 0.3%-Triton PBS and primary (ON at 4°C) and secondary antibodies (3h at RT) were used in 0.1%BSA PBS and are listed in Supplementary Table 2. After staining, tissue-slides were mounted to a microscope slide using Vectashield. The samples were imaged using a SP8 confocal microscope (Leica Microsystems, 40x oil objective, N.A. = 1.3).

### Organoid culture

All organoids were obtained from the duodenum of male or virgin female mice older than 8 weeks as described previously^52^. Organoids were embedded in BME and cultured with ENR media including Advanced DMEM/F-12 (Life Technologies) supplemented with; murine recombinant epidermal growth factor (50 ng/ml; Life Technologies), murine recombinant Noggin (100 ng/ml; PeproTech), human recombinant R-Spondin 1 (500 ng/ml; Peprotech), n-acetylcysteine (1 mM; Sigma-Aldrich), N2 supplement (1×; Life Technologies) and B27 supplement (1×; Life Technologies), GlutaMAX (2 mM; Life Technologies), Hepes (10 mM; Life Technologies), and penicillin-streptomycin (100 U/ml; 100 μg/ml; Life Technologies). For passaging, organoids were removed from BME and mechanically dissociated into single-crypt domains, and then transferred to fresh BME. Passage was performed every 1-2 weeks with a 1:5 split ratio. Conditions were kept stable in the incubator with 5% CO2 and 20% O2 at 37°C.

### 2D organoid culture

#### Hydrogel preparation

For the 2D organoid cultures, a hydrogel composed of a collagen mixture was used to mimic the extracellular matrix (ECM). All materials and reagents, including Eppendorf tubes, stock collagen (5 mg/mL Atelocollagen, ReproCELL, cat no. KKN-IPC-50), buffers, and Matrigel, were pre-chilled on ice to prevent unwanted polymerization of the precursor gel due to prolonged exposure to heat.

A collagen gel precursor (4 mg/mL) was prepared by neutralizing the pH of the collagen stock with basic R-buffer and 10x DMEM in an 8:1:1 volume ratio. The precursor was mixed by slow and gentle pipetting to avoid introducing air bubbles. The phenol red in the 10x DMEM changed from yellow to light pink, indicating the shift in pH of the gel precursor from acidic to basic. Due to the high viscosity of the gel precursor solution, volumes were kept small, for example, 160 µL collagen stock, 20 µL R-buffer, and 20 µL 10x DMEM.

The final hydrogel mixture was prepared by combining the gel precursor with Matrigel in a 4:1 volume ratio, resulting in a final concentration of 3.2 mg/mL. The mixture was homogenized by slow and gentle pipetting.

To polymerize the hydrogel mixture, a LabTek 8-well chambered coverglass (Thermo Fisher Scientific, cat. no. 155411PK) was pre-treated with plasma treatment. Prior to gel transfer, the coverglass was pre-chilled on ice, preferably on a metal heating block for even heat transfer, to prevent unwanted polymerization. 17.5 µL of the hydrogel mixture was transferred to each chamber, ensuring even coverage of the entire bottom surface. The coverglass was then immediately placed into a 37°C incubator and kept for 30-min until polymerized.

#### Seeding

The 3D intestinal organoids were mechanically disaggregated by pipetting in cold Advanced DMEM/F-12 (Life Technologies). For one chamber in the LabTek 8-well chambered coverglass, we seeded approximately 0.7x the number of organoids contained in a Matrigel drop of the 24-well plate. After disaggregation, the organoids were centrifuged at 300g for 5 min at 4°C and the pellet was resuspended in ENR medium. The organoids were seeded in a small volume (150 µL) in each chamber and incubated at 37 °C and 5% CO2. ENR medium was refreshed every 2 days. All experiments were performed 4 d after seeding.

### Microfluidic chip preparation and experiment

Microfluidics experiments were performed as described previously^38,53^. In short, a mold for microfluidic chips was made using a 3D printer (Atum3D) (chip design available upon request). PDMS (SYLGARD #01673921/ #01673921, 1:8 ratio) was poured in the mold to create the microfluidic chip. The chip and a glass slide (Marienfeld, No 1.5H, high precision, 70×70 mm) were bonded using plasma treatment (Diener electronic, Femto). Chip and Tubing were UV cleaned and the microfluidic chip (in PBS with Pen/strep) was degassed together with the conditioned and DAPT media for 4 hours. The concentration of the DAPT media depended on the pulse timing (90min: 1.38µM, 130min: 2µM, 170min; 2.6µM, 2µM; for 170 min we tested two different concentrations to find one that did not change oscillation amplitude or absolute intensity (Fig. 4 and Extended Data Fig. 12). Organoids were passaged 3-4 days before performing a live-imaged microfluidic experiment for 20-24h. For the RNA bulk sequencing and immunofluorescence microfluidic experiments organoids were passaged 1 day before a 72h experiment, 2 days before 48h experiment and 3 days before a 24h experiment. The day of the experiment, two wells of a 24-well plate were harvested and washed with ice cold Advanced DMEM/F12 supplemented with GlutaMAX (2 mM; Life Technologies), Hepes (10 mM; Life Technologies), and penicillin-streptomycin (100 U/ml; 100 μg/ml; Life Technologies) to remove the BME. Organoids are resuspended in BME to a final concentration of 50% and carefully loaded into the microfluidic chip which was pre-filled with ENR culture media. After loading organoids on the chip, all tubing (4x inlet tubing of 3 meters, and 2x outlet tubing of 1 meter) was connected and during the experiment the organoids were cultured under a flow rate of 200 µl/h, using a microfluidic pump (Harvard apparatus #70-45060). After approximately 30 min culturing in culture medium, pumping program to induce drug pulses was started. For the bulk RNAseq and immunofluorescence microfluidic experiments, organoids were removed by aspirating the organoids out of the chip with a 200ul pipet via the organoid inlets and follow-up methods were performed as described.

### Time-lapse imaging

#### Intravital imaging

Intravital imaging was performed as described in detail in Laskaris et al., 2023^54^. In short: Time-lapse intravital imaging was performed on the healthy gut of Hes1-Achilles reporter mice in H2B-mCherry background. Throughout the procedure, mice were sedated using isoflurane inhalation anaesthesia (1.5% isoflurane/ O2 mixture). The gut was surgically exposed, and the mouse was placed in a custom designed imaging box with its head in a facemask constantly delivering anaesthesia. The imaging box and the microscope were adjusted at 36.5°C using a climate chamber. Intravital images were acquired with an inverted Leica TCS SP8 AOBS two-photon microscope (Mannheim, Germany) with a chameleon Ti:Sapphire pumped Optical Parametric Oscillator (Coherent Inc. Santa Clare, CA, USA). The microscopes are equipped with 2 non-descanned and 2 hybrid detectors: YFP and RFP were simultaneously excited at 960 nm and detected with hybrid detectors. Second harmonic generation (Collagen I stroma) was detected with a non-descanned detector. Images were collected every 10 min for a period of 3-6 hours during which the mouse was kept sedated and alive, constantly hydrated with subcutaneous infusion of glucose and electrolytes (NutriFlex special 70/240, Braun, 100 μl/h). All images were collected in 12 bit, 256X256 and acquired with a 25X water immersion (HC FLUOTAR L N.A. 0.95 W VISIR 0.17 FWD 2.4 mm) objective and zoom of 1.25X.

#### SP8 imaging of 3D organoids

For Hes1-Achilles intensity analysis of crypts, organoids were passaged 3 days before every live imaging experiment on an 8 chambered Lab-Tek Borosilicate microscope dish with a cover glass (Thermo Fisher Scientific, cat. no. 155411PK). The organoids were imaged in fresh ENR and drugs diluted in ENR were added after imaging the first time point or at a later timepoint when indicated. The Leica SP8 confocal microscope was used for live imaging of the organoids and were present a humid atmosphere at 5% CO2, 20% O2 and 37°C. The fluorescent Achilles protein was excited using a 514 nm laser and organoids were visualized using the 20x objective. Every 30 min images were taken, using the bidirectional scanning function and a frame average of 4. To visualize the full organoid, multiple images were taken from each organoid spaced by z-steps of 6 µm and a pinhole of 184.5 μm (PinholeAiry of 3.26).

#### 3D *organoid Light-sheet imaging*

Live imaging of organoids for single cell analysis was performed using a Viventis light sheet microscope (Viventis microscopy #LS1 live). Imaging was performed in a humid atmosphere at 5% CO2, 20% O2 and 37°C. 514, 568, 687 nm lasers were used for Achilles, mCherry and iRFP, respectively and images were taken with a 25x objective. A Laser beam of 2,2 µm was used and a z-stack of 100 steps of each 2 µm was taken. Images were taken at 1024 x 1024 pixels every 7 min.

#### Organoids in microfluidic chip

Live imaging of organoids in the microfluidic chip was performed using a Leica SP8 MP or SP5 MP confocal microscope in a humid atmosphere at 5% CO2, 20% O2 and 37°C. Organoids were excited with a 514nm laser for Achilles through a 20x objective. Every 10 min a z stack of 8µM distance was scanned at 512×512. To track given drug pulses Cascade blue (cat. No. Cat#C3239) was added to the DMSO or DAPT supplemented media and excited with a multi photon laser at 740nm and 3x 30µM z planes were detected every 10 min at 64×64 pixels.

#### 2D *organoids*

Four days after passaging, 2D organoids were imaged in an 8-well chambered coverglass using confocal microscopy (Leica SP8) with a 25x objective. Environmental conditions were set at 5% CO2, 20% O2, and 37°C with high humidity. Samples were placed inside the microscope and incubated for over 2 hours before imaging to allow for acclimatization. Hes1-Achilles was excited at 514 nm and H2B-mCherry at 564 nm using a diode laser. Timelapse images were captured every 7 min with a resolution of 512×512 pixels at zoom 1 (pixel size: 0.36 µm), and z-steps of 0.4 µm, using a pinhole of 1 Airy unit.

### Immunofluorescence of 3D-cultured organoids

Organoids in BME were fixed with 4% paraformaldehyde (freshly made at same day) for 10-45 min at RT (Supplementary Table 2), washed 3 times with PBS and blocked in PBS containing 5% Normal Goat Serum (NGS) and 0.3% Triton X-100. Samples were incubated with the primary antibodies in 1% BSA, 0.3% Triton-X100 in PBS (Supplementary Table 2) ON at 4 °C while shaking, after which the samples were washed 3 times with PBS for 10 min. The samples were incubated with Alexa conjugated secondary antibodies (1:500; Life Technologies) and DAPI (5 μg/mL, Sigma) for 3h at room temperature while shaking and subsequently washed 3 times with PBS for 10 min. The samples were imaged using a SP8 confocal microscope (Leica Microsystems, 40x oil objective, N.A. = 1.3).

### Immunofluorescence of 2D-cultured organoids

2D organoids were fixed in 4% paraformaldehyde (PFA) for 20 min at room temperature (RT) and washed three times with phosphate-buffered saline (PBS). The samples were permeabilized with 0.1% Triton X-100 (Sigma-Aldrich) for 45 min at RT. After three washes with PBS, the samples were blocked with goat serum for 1 hour at RT. Primary antibodies diluted in PBS containing goat serum were added and incubated overnight at 4°C. Following three washes with PBS, secondary antibodies in PBS containing 10% goat serum were added and incubated for 2 hours at RT. After three additional washes in PBS, wheat germ agglutinin (WGA) diluted to 1:500 in PBS was added for 1 hour at RT. Finally, the samples were washed three times with PBS (5 min each) and imaged.

### Image and data processing

Gut-rolls were analysed using Fiji and graphs were plotted with Prism.

Microfluidic time series were analysed using a Fiji Macro, based on manual tracking of a ROI and plotted in Python. Quantification of crypt oscillatory dynamics was performed by using PyBOAT (settings: min_period=50, max_period=400, TC=200).

Kymographs were created as follows: First, a maximum intensity projection was performed on all z-planes. The time series data were then smoothed using a Gaussian filter (2 scaled units). Kymographs were constructed along a user-defined line extending from the crypt to the villus using the Fiji^55^ plugin KymoResliceWide (intensity averaged over the width of a 50 px wide line). These kymographs were further smoothed using a Gaussian filter with 2 scaled units.

Intravital imaging images were manually corrected in XY using a Fiji plug-in Bigdataprocessor2. Z correction and single-cell tracking was manually performed in Mastodon a Fiji plug-in. Single-cell tracks were plotted in phyton.

3D organoid light-sheet images were cropped and registered using LStree^56^. Single cells were manually tracked using Mastodon. Hes1-Achilles intensity was normalized to the global mean of Hes1-Achilles per experiment. Graphs were plotted with Python and quantification of oscillation dynamics was performed using pyBOAT (settings: min_period=50, max_period=400, TC=200) and Python.

Image Analysis of stainings and markers after microfluidics and average DAPT included (Fig. 5 & Extended Data Fig. 16, 17):

#### 1. Organoid segmentation

Image analysis was performed using Python 3 and Scikit-image [scikit-image 0.24.0 (2024-06-18)]. Organoids were segmented from each acquired z-stack based on the DAPI channel. For each plane, a mask was generated by applying a Gaussian blur and binarization with a threshold that was set at 2 times the mean DAPI intensity of the plane. Following binarization, masks with fewer than 2000 pixels were excluded from the analysis. All pixels outside the mask were set to zero. Maximum intensity projections (MIPs) were generated from all planes and used for further analysis. For each segmented organoid, the area and the mean intensity of markers was calculated.

#### 2. Superpixel analysis on organoid masks

Superpixel analysis was conducted following the protocol established by Suppinger et al. (2023)^57^. The DAPI mask in each image was considered as the total superpixel value, with each superpixel approximating the size of a single cell, approximately 250 pixels. Superpixels served as proxies for individual cells to quantify the presence of LGR5, WGA, SOX9, LYZ1, and SCA-1 expressing cells. The mean intensity of the markers in superpixels in control organoids (DMSO) was used as a threshold to identify positive superpixels in the test condition (DAPT-treated). The percentage of positive superpixels was then calculated relative to the total number of superpixels per cell type marker. Additionally, enterocyte percentages were inferred by subtracting LGR5+ and WGA+ [and SOX9+] superpixels from the total number. To ensure that our analysis was focused on homeostasis, all organoids smaller than 100000 pixels were excluded from the analysis. This size exclusion threshold was based on manual selection.

##### 2D *image analysis*

###### Segmentation and tracking

Nuclear annotation was performed using Cellpose with the “nuclei” model as a starting point. A total of 306 images was annotated in each xy, xz, and yz planes. The annotations generated by Cellpose served as ground truths to train Stardist, ensuring star-convex shapes for nuclei.

Cell tracking was conducted using the Laptrack algorithm in Python. Tracks were manually verifying in high-division zones near the crypt centres, addressing segmentation errors that often occurred during cell division. The Laptrack-generated tracks were converted to GraphML files and manually verified using Mastodon, a plugin for Fiji.

###### Cell Type Classification

To classify cell types by backtracking from endpoint stainings, the last frame of live imaging was manually aligned to endpoint IF stainings. Cells in this frame were manually annotated per cell type in Napari software in Python. Custom Python code allowed cell type back-tracking with the following rules:

1. Stem cells are OLMF4+ and the mother cells of stem cells are stem cells, assuming the directionality of differentiation.
2. Secretory cells are WGA+
3. Paneth cells are WGA+, adjacent to stem cells and do not divide ^35^ during the 24-hour imaging period

Using these definitions, we calculated the distance from the crypt centre as the number of cells away from Paneth cells (Fig. 2)

4. IMPCs are mothers of secretory cells or mothers of Hes1-Achilles low cells within the crypt (<4 cells aways from Paneth cells) which lacked the WGA marker.

We assumed that these WGA-negative cells are in fact IMPCs that have committed to their fate but have not yet produced mucin detectable by WGA. Or they belong to a rare secretory lineage such as enteroendocrine cells. However, given the statistical improbability of their being rare cell types, we classified them as IMPCs.

5. Enterocytes are Hes1-Achilles low cells (track intensity median below threshold = 0.075), 2) WGA negative and 3) far away (>3 cells) from Paneth cells.

Based on our staining for enterocytes in 2D cultures (Extended Data Fig. 6), we know that enterocytes make up of most of the non-secretory, Hes1-Achilles low cells residing on the outside of the crypt zone.

6. TA cells are Hes1-Achilles positive cells residing between stem cells and enterocytes

a. Early TA are Hes1-Achilles high and up to 3 cells away from Paneth cells
b. Late TA are Hes1-Achilles high and 4 cells or more away from Paneth cells.

###### Hes1-Achilles Intensity Track Analysis

All analysis was performed using custom-written Python code. To standardize Hes1-Achilles intensities across datasets, intensities were normalized to the minimum and maximum values of each dataset. Tracks were smoothed with a moving mean across three time points to remove non-biological noise.

Fourier transform was performed for each track, and then the fundamental and the first harmonic contributions were removed for each track to isolate genuine oscillatory signals. Mean power spectrum was generated across all tracks. Wavelet analysis was implemented using PyBOAT, with a minimum and maximum period (Tc) set to 200. The period of best fit was calculated using ridge detection within the PyBOAT algorithm. Neighbourhood analysis was adapted from methods available on Robert Haase’s GitHub repository. Calculation of the Kuramoto order parameter was adapted from the Arnold-Tongue paper.

To detect transition events, we selected a subset of tracks that had low Hes1-Achilles intensities (<0.075 AU) in the last 20 frames (140 min). We then applied a rolling median of 10 timepoints (70 min) to Hes1-Achilles reporter intensities in each track to reduce noise. We defined the time when this median first becomes >0.1 AU as the transition point.

### Statistical Analysis

For all experiments at least three independent experiments were performed. To visualize data, Tukey style boxplots were used (bar=median, box=25th and 75th percentile, whiskers=1.5*IQR). Data displayed as violin plots present data from max to min with bar=median. For statistical comparisons between groups, normal distribution was calculated by Shapiro-Wilk testing. Significance was tested by independent t-test or Mann-Whitney U test depending on normal distribution. To correct for multiple groups Bonferroni calculation was used.

### RNAseq quantification and Statistical Analysis

Organoids (20-50 organoids per sample) were harvested from the microfluidic chip. Total RNA was isolated using TRIzol Reagent (Invitrogen, #15596018). In brief, samples were barcoded with CEL-seq2 primers during a reverse transcription and pooled after second strand synthesis. The resulting cDNA was amplified with an overnight *in vitro* transcription reaction. From this amplified RNA, sequencing libraries were prepared with Illumina Truseq small RNA-primers. The DNA library was paired-end sequenced on an Illumina NextSeq™ 500, high output, with a 1×75 bp Illumina kit (R1: 26 cycles, index read: 6 cycles, R2: 60 cycles). Reads that mapped equally well to multiple locations were discarded. Mapping and generation of count tables was done using the STARSolo 2.7.10b aligner.

### Bulk RNAseq mapping and analysis

Resulting bulk RNAseq data were mapped to the mouse genome (mm38) using STAR (version 2.7.8a). umi_tools (version 1.1.1) was used to collapse PCR-amplified reads into distinct transcripts. Samples with less than 500,000 transcripts were determined as low quality and discarded from further analysis, resulting in median depth of 960,086 transcripts per sample. To generate heatmaps (Figure 5b, Extended Data Figures 15a-b), data was size normalized, and each sample was compared to the time-matched DMSO controls.

### Reanalysis of single-cell RNAseq data

Single-cell RNAseq data from Bues et al. (2022)^40^ were downloaded and re-clustered with the Metacell package, as described before^58^. Cells with less than 400 UMI or more than 20,000 UMI were treated as low quality or doublets and discarded from analysis, resulting in 2,285 total cells retained for clustering analysis.

We selected genes as features for the metacell algorithm using the parameters Tvm=0.06, total umi > 5, and more than 4 UMI in at least 3 cells. We filtered the list of gene features from genes associated with cell cycle or immediate stress response. For this, we first identified all genes with a correlation coefficient of at least 0.12 for one of the anchor genes *Top2a*, *Ube2c*, *Mki67*, *Pcna*, *Mcm4*, *Egr1*, *Fos*, and *Jun*. We then hierarchically clustered the correlation matrix between these genes and selected the gene clusters containing the above anchor genes. We thus retained 153 genes as features. We used Metacell to build a kNN graph, perform boot-strapped co-clustering (500 iterations; resampling 70% of the cells in each iteration), and derive a cover of the co-clustering kNN graph (K=20). We manually split meta-cells that mixed goblet and Paneth cell signatures by clustering these cells based on gene modules correlated with *Muc2* (goblet cells) and *Lyz1* (Paneth cells).

scRNAseq Data from Haber et al. (2017)^12^ were downloaded and harmonized with the Bues et al. dataset as follows: For each single cell in Haber et al. (representing wildtype organoids grown in ENR only), we computed the K (K = 10) nearest neighbors from Bues et al. based on Pearson correlation. Then meta-cell assignment for each cell in Haber et al. was determined by majority voting.

### LDA-based bulk RNAseq deconvolution

We applied a Latent Dirichlet Allocation (LDA)-based approach to predict cell type distribution of samples in the bulk RNAseq dataset^59^. Whereas in conventional LDA analysis, both the distribution of topics as well as the probabilities of observing words within topics are determined concurrently, here, we used predetermined topics extracted from analysis of a reference scRNAseq dataset. Thus, each metacell in the reanalysis of Bues et al. represents a topic, whose words (i.e., genes) distributions is calculated as a multinomial probability distribution. We use the LDA algorithm (topicmodels version 0.2-9) with a predetermined beta matrix, in order to optimize the distribution of topics per text (i.e., metacells per bulk sample). We selected a subset of the genes and used them as features for the LDA as follows: We selected for each metacell the top 30 most enriched genes, given they have at least 0.7 enrichment score (log_2_ fold-change over the median expression), and at least 100 UMI across all single cells and at least 1,000 transcripts in the bulk dataset. This resulted in a set of 342 genes used as features. To increase reproducibility, we reran the LDA 100 times, each time with a slightly altered beta matrix. The beta matrix (representing the words per topic multinomial distributions) in each iteration was calculated over a sample of the single cells (representing 70% of cells per meta-cell). We then extracted the median value per meta-cell.

We sought to test the performance of this approach across two unrelated datasets. For this purpose, we tested our ability to deconvolve pseudo-bulk aggregates generated from Haber et al. (generated with the 10x genomics platform), using Bues et al. (generated by the DisCo technology) as a single-cell RNAseq reference. We first harmonized the two datasets as described above. Then, we generated 100 pseudo-bulk aggregates of 200 cells from Haber et al., mixing cell types at varying proportions. We used our LDA approach to infer cell type proportions for each aggregate and compared them to the ground truth. We report high agreement between inferred and ground-truth levels of Enterocytes (R^2^ 0.85), Enteroendocrine cells (0.78), Goblet cells (0.72) and Paneth cells (0.88), whereas stem cells and TA cell levels could not be faithfully inferred (R^2^=0.17 and 0.15, respectively), probably due to them lacking a clear transcriptional signature.

## Data and Code availability

All data and custom code used in this study are available upon request to.

## Acknowledgements

We thank all members of the Sonnen, Bothma and Garaycoechea labs for discussions and Juan Garaycoechea and Takashi Hiiragi for feedback on the manuscript. We thank Ben Simons, François Schweisguth, Johan van Es and Hans Clevers for insightful discussions, Joep Beumer for advice when setting up organoid culture in the lab, Jean-Yves Tinevez for developing a beta-version of Mastodon, Mike Nikolaev, Nikolce Gjorevski and Matthias Lütolf for support with setting up the 2D organoid culture and Yasmine el Azhar for providing mice. We thank Wouter Thomas, Jeroen Korving, Benaissa el Haddouti, Stieneke van den Brink and Maria Azkanaz for technical help. We are grateful to Jos Malda for sharing microfabrication equipment and Andrea Boni (Viventis) for advice. This work was supported by the Hubrecht imaging and animal facilities and the NKI animal facility. The H2B-mCherry was kindly provided by the Riken Centre for Biosystems Dynamics Research, the Lgr5tm2(DTR/EGFP)Fjs mouse line by Fred de Sauvage, Genentech, and the iRFPnucmem reporter by Alexander Aulehla. This work was supported by the Hubrecht Institute and received funding from the European Research Council under an ERC starting grant agreement no. 850554 and KWF grants 13661 and 15261 to K.F.S.

## Author contributions

Conceptualization: S.W., H.E. and K.F.S., Methodology: S.W., H.E., K.A, E.L., M.H., W.B., Software: H.E., J.L. and A.G. Formal Analysis: S.W., H.E., J.L., A.G. and K.F.S., Investigation: S.W., H.E., Writing: S.W., H.E. and K.F.S., Review and Editing: S.W., H.E., J.L., A.G., J.R., A.O. and K.F.S., Supervision: J.R., A.O. and K.F.S., Project administration and Funding acquisition: K.F.S.

## Supplementary Information

### Extended Data Fig. Legends

**Extended Data Fig. 1.**
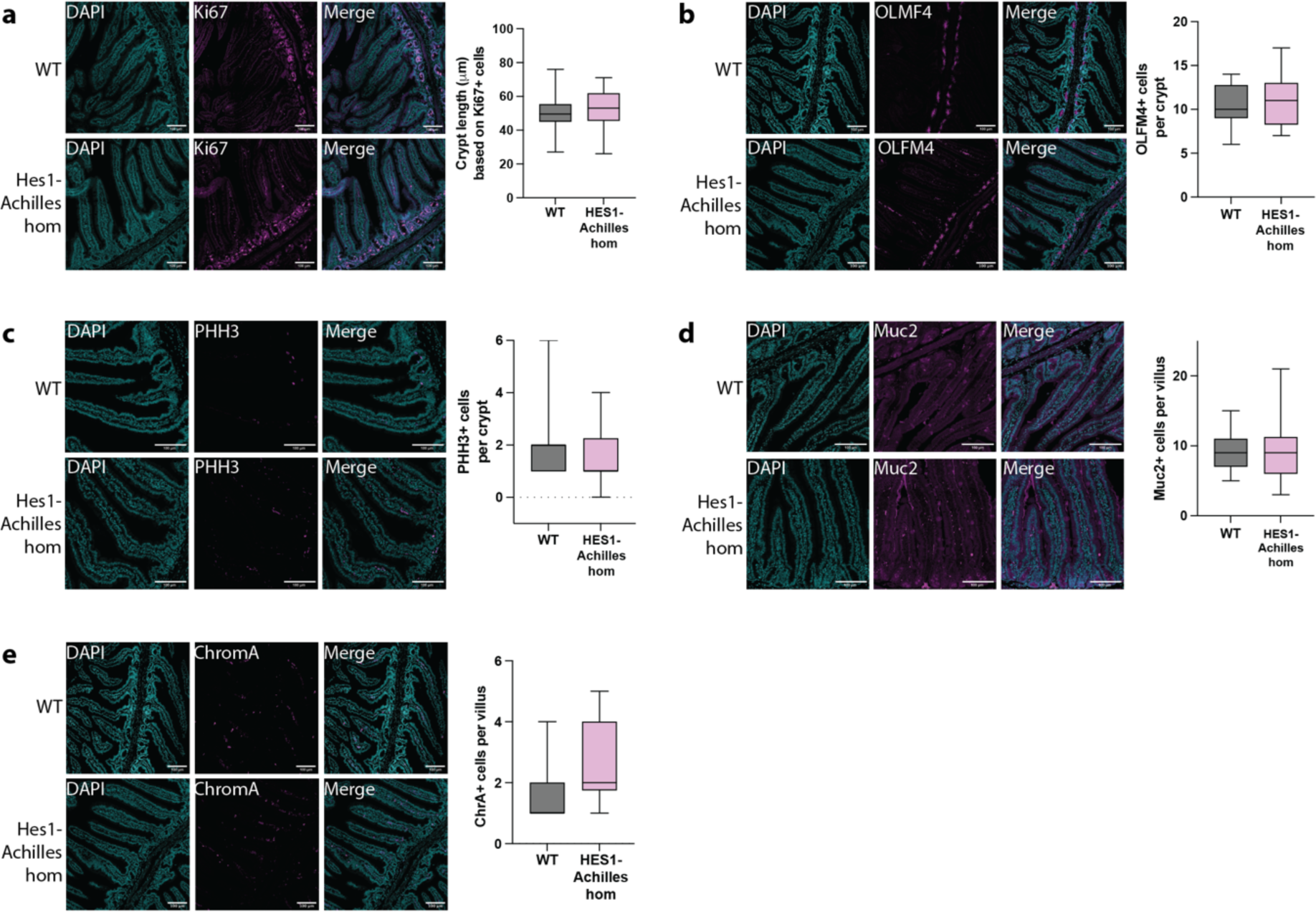
Characterization of the small intestine in Hes1-Achilles reporter mice. Representative images of WT and Hes1-Achilles gut-rolls immuno-stained for a proliferative marker Ki67 (a) or TA cell marker OLMF4 (b) or mitotic marker PHH3 (c) or goblet cell marker Muc2 (d), or enteroendocrine cell marker ChrA (e) and counterstained with DAPI. The length (a), amount per crypt (b, c) or amount per villus (d, e) of cell marker signal was quantified in WT and Hes1-Achilles gut rolls. 2 male and 2 female mice were used for both conditions and 3 crypts or villi were used to sample the cells or crypt size. No significant differences were detected. Normal distribution was calculated by Shapiro Wilk test and significance was tested by independent t-test or Mann-Whitney U test depending on normal distribution. ns: 5.00e-02 < p <= 1.00e+00, *: 1.00e-02 < p <= 5.00e-02, **: 1.00e-03 < p <= 1.00e-02, ***: 1.00e-04 < p <= 1.00e-03, ****: p <= 1.00e-04.

**Extended Data Fig. 2.**
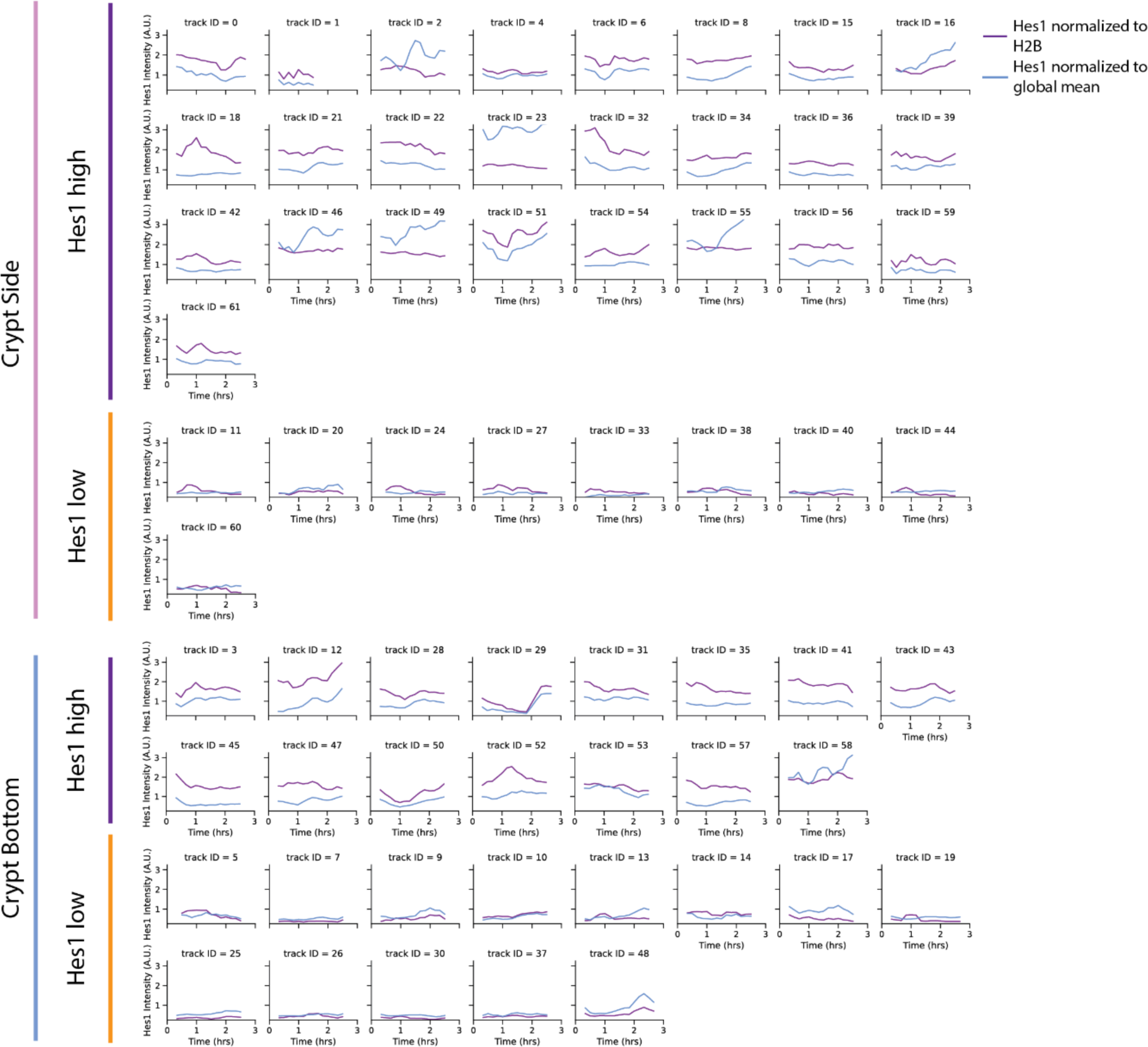
Single-cell tracks measured in small intestinal crypts *in vivo*. Hes1-Achilles mean-normalised (blue) and Hes1-Achilles H2B normalized (purple) timeseries data of single cells obtained by intravital imaging of the small intestine (N=1, n_crypt=4). Tracks have been manually subdivided into crypt bottom and side (total n=62 tracks), and clustered on Hes1-Achilles low (Hes1-Achilles H2B normalized <0.8) and Hes1-Achilles high (Hes1-Achilles H2B normalized >0.8).

**Extended Data Fig. 3.**
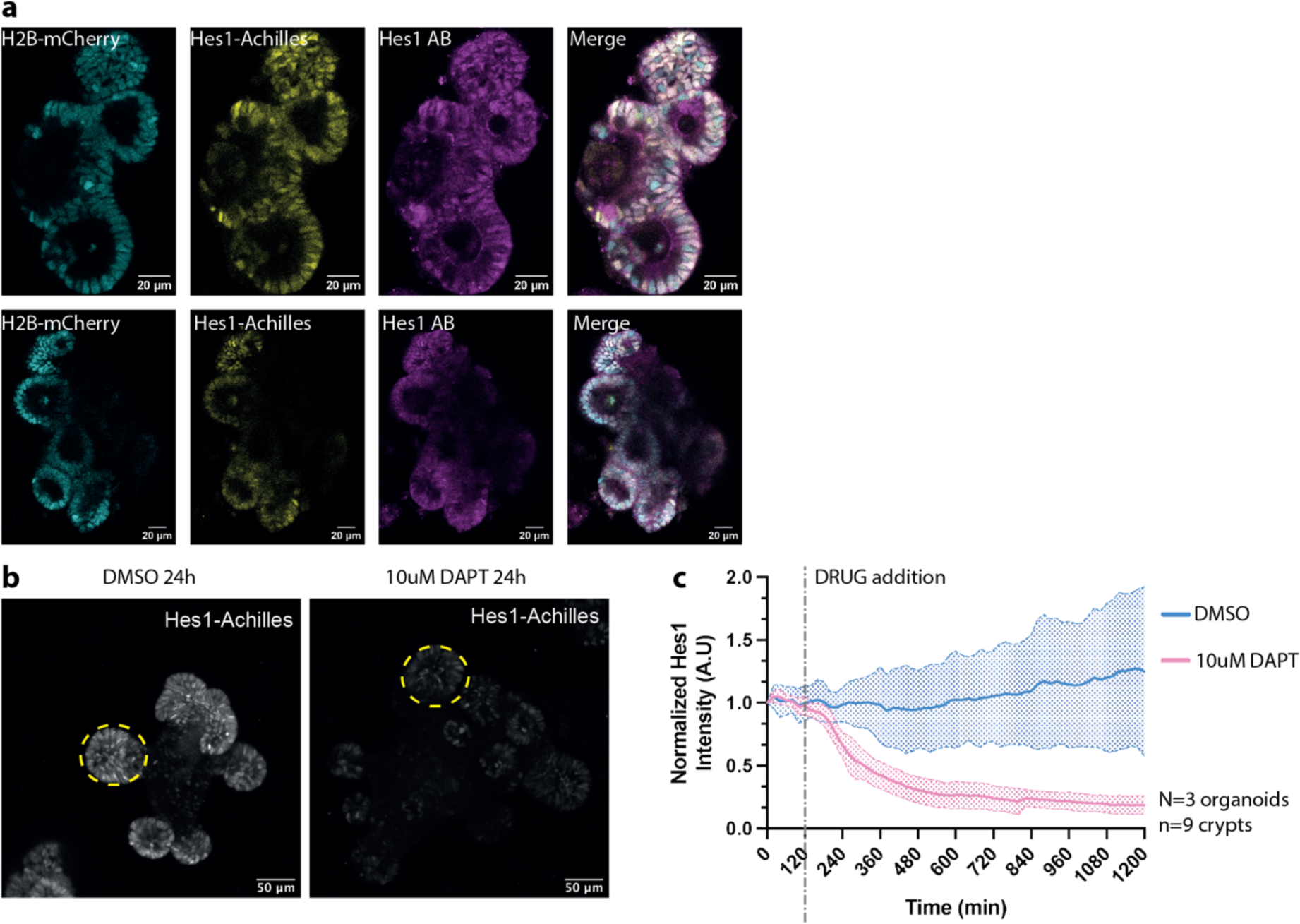
Hes1 expression is NOTCH dependent in intestinal organoids. (a) Representative images of Hes1-Achilles small intestinal organoids, with nuclear marker H2B-mCherry, immuno-stained for Hes1. (b) Representative images of small intestinal organoids treated with DMSO (control) or the gamma-secretase inhibitor DAPT after 24 hours. (c) Hes1-Achilles intensity (normalised to first 120 timepoints of DMSO) was measured over time. Dotted line represents the SD and solid line represents the mean of 9 crypts tracked in 3 organoids. Grey line represents treatment start time of 120 min.

**Extended Data Fig. 4.**
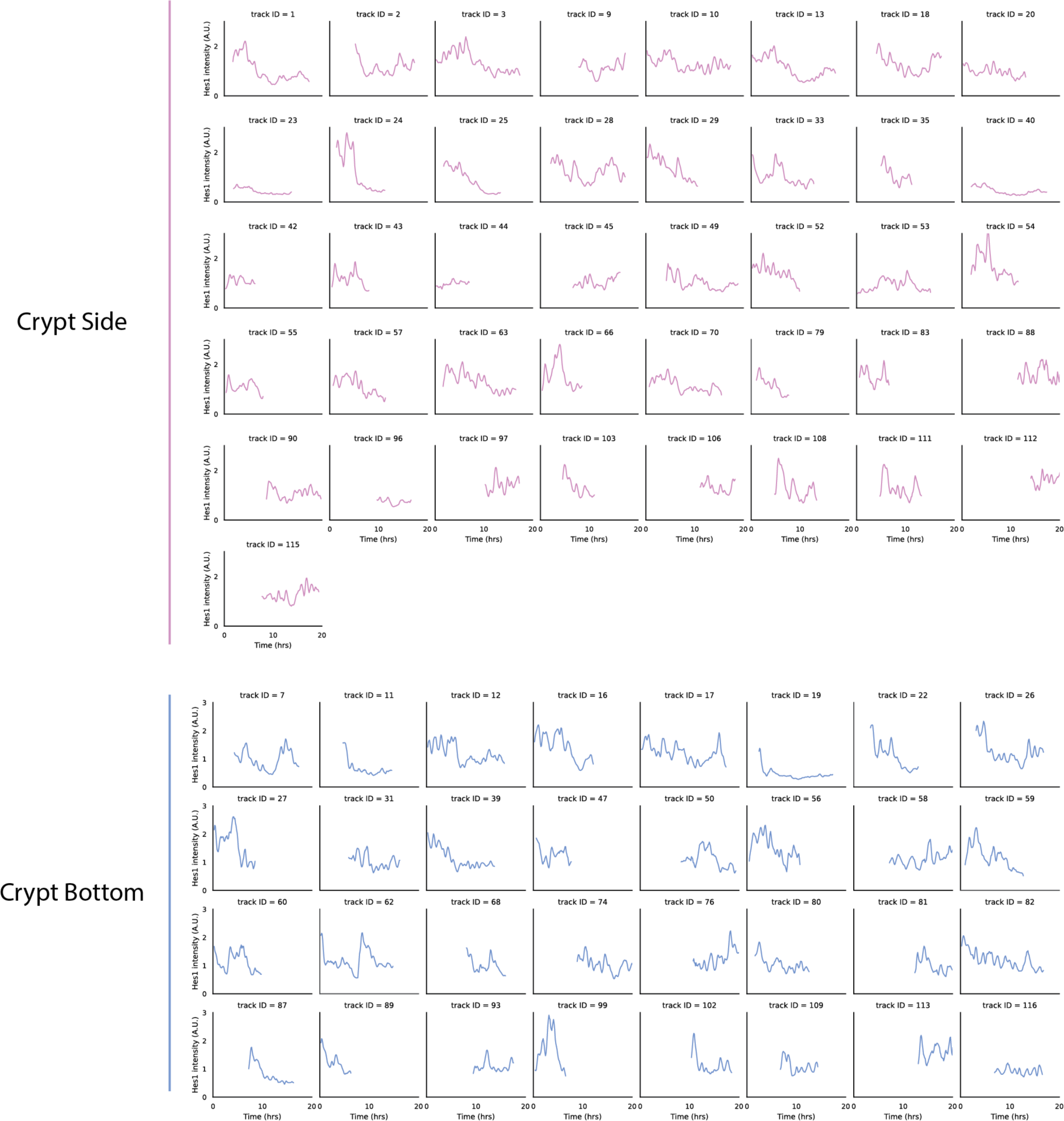
Tracks of single cells in crypts of Hes1-Achilles intestinal organoids. Mean-normalised and detrended timeseries data of single cells analysed in Fig. 1i, j (N=3). Single-cell tracks are manually subdivided into crypt bottom (n=32) and side cells (n=41).

**Extended Data Fig. 5.**
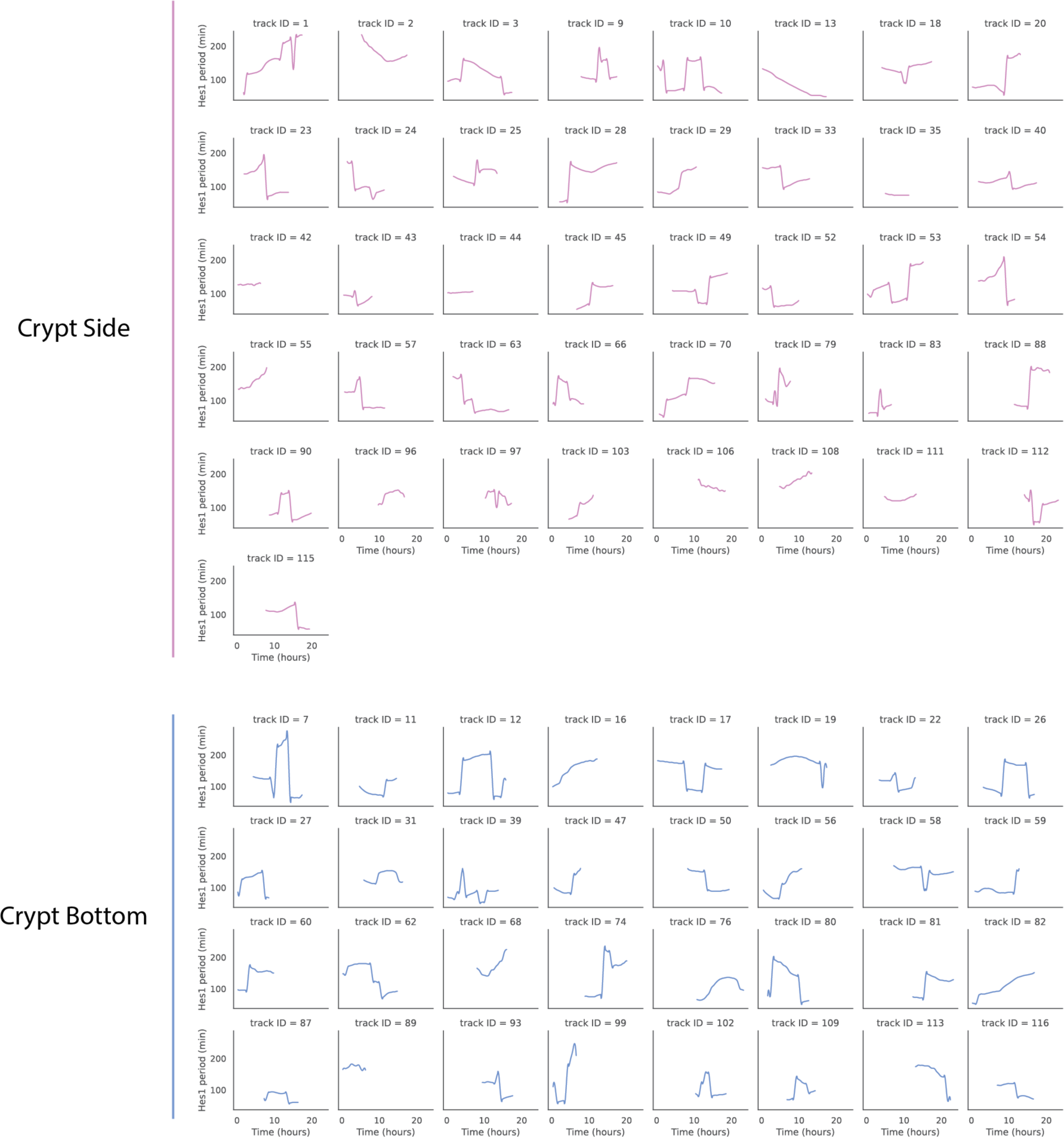
Instantaneous period of single cells in crypts of Hes1-Achilles intestinal organoids. Period quantified by wavelet transform over time which is plotted in Fig. 1k (N=3). Single cell period tracks are subdivided into crypt bottom (n=32) and side cells (n=41).

**Extended Data Fig. 6.**
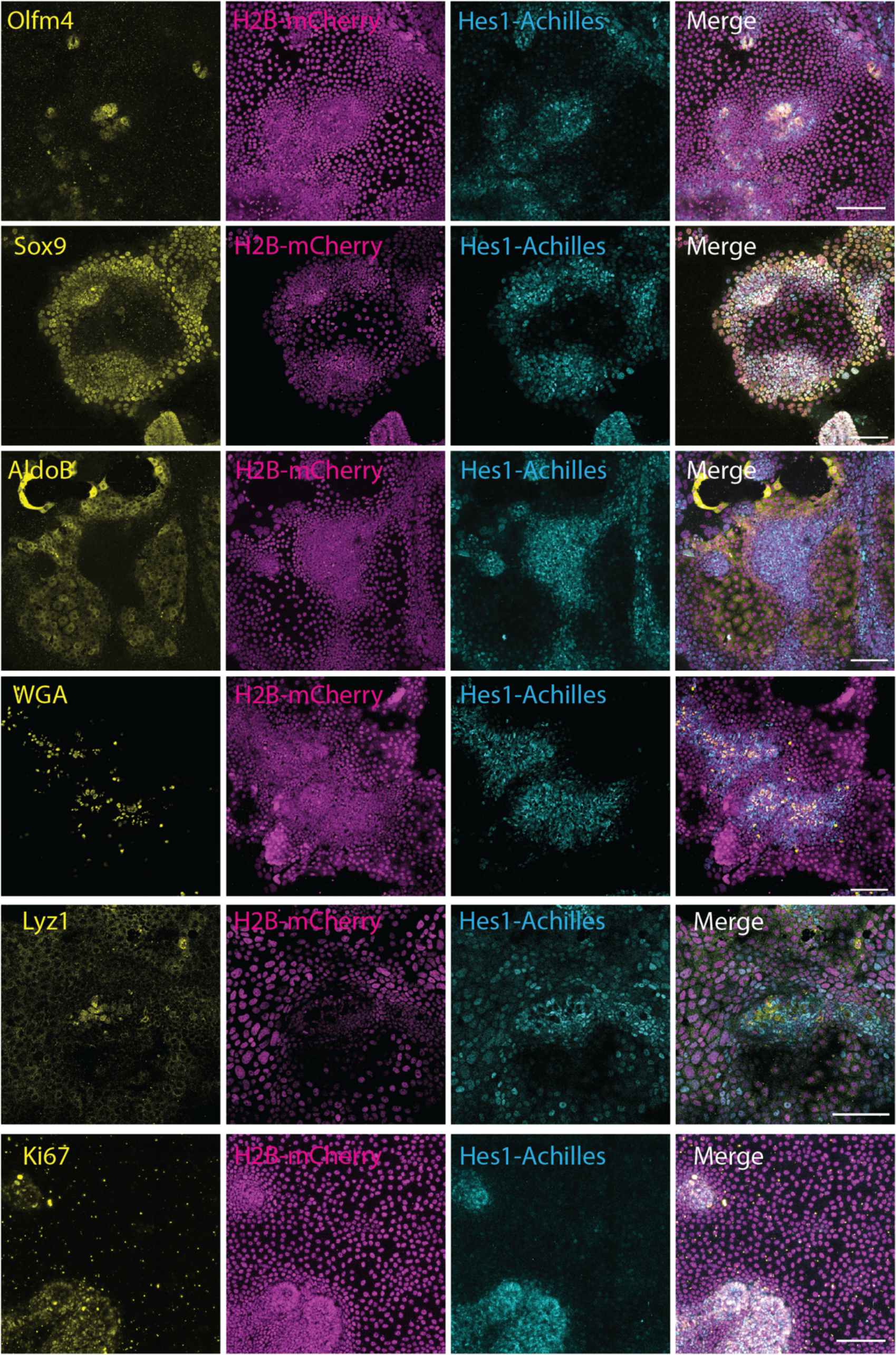
Immunofluorescent staining of cell-type markers. Intestinal organoids expressing Hes1-Achilles and H2B-mCherry were cultured in 2D, followed by IF stainings for the indicated markers as well as WGA staining. Representative images are shown. Scale bar 50 µm.

**Extended Data Fig. 7.**
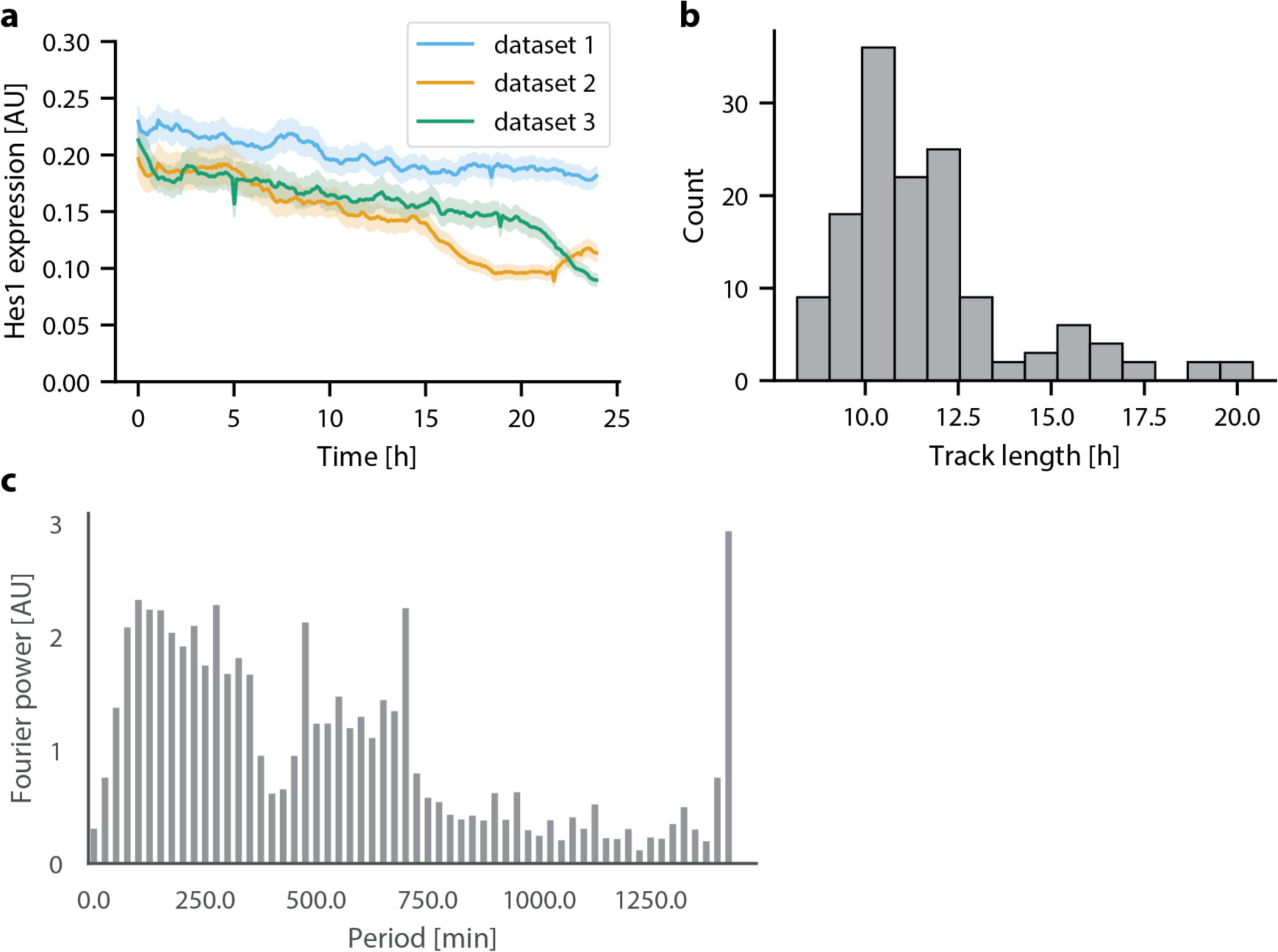
Further analysis of single-cell Hes1-Achilles traces. Intestinal organoids expressing Hes1-Achilles and H2B-mCherry were cultured in 2D, followed by live-imaging and analysis. (a) Plot of Hes1-Achilles intensity vs time for all single-cell tracks, track n_dataset1_ = 2729, n_dataset2_ = 2102, n_dataset3_ = 1705. (b) Histogram of track length from division to division, for all proliferative cells, n = 140. (c) Mean Fourier power spectrum for all single-cell Hes1-Achilles intensity traces, before removing contributions from harmonics related to track length (after removal in Fig. 2b). N=3 biological replicates.

**Extended Data Fig. 8.**
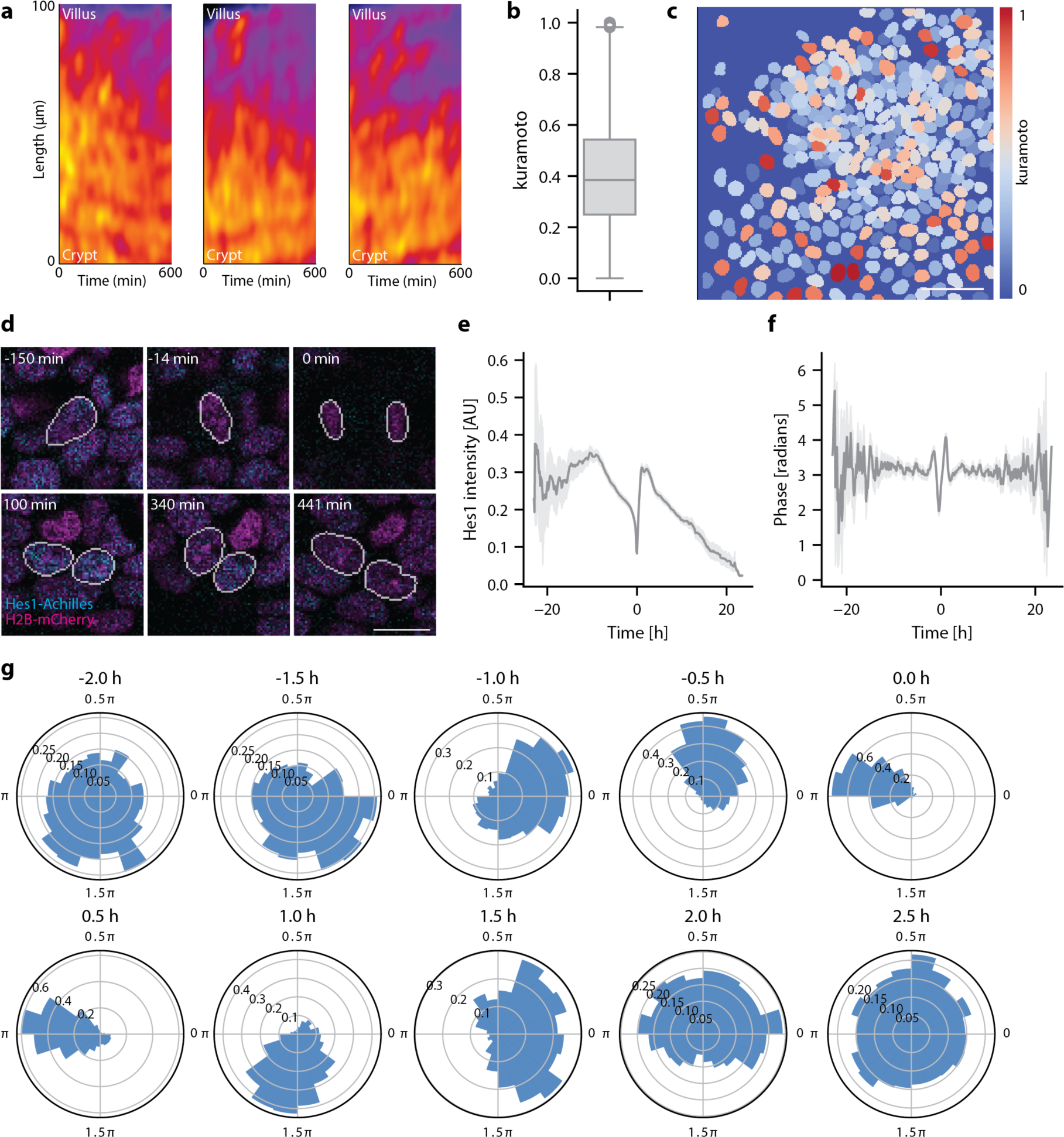
Local synchronisation of Hes1 dynamics. Intestinal organoids expressing Hes1-Achilles and H2B-mCherry were cultured in 2D, followed by live-imaging and analysis. (a) Examples of kymographs taken from timelapse images lines of 2D organoids expressing Hes1-Achilles (see Fig. 2d). (c) Boxplot of Kuramoto order parameter calculated for the direct neighbours of all cells at all timepoints, n_inst_ = 147959 instantaneous measurements. (d) Representative snapshot of segmented nuclei showing Kuramoto order parameter. Scale bar: 50 µm. (e) Representative snapshots of a dividing cell, taken from live imaging of 2D organoids. Scale bar: 5 µm. (f) Hes1-Achilles oscillation phase and (g) intensity vs time for all dividing cells, centred around division, n = 905 tracks. (h) Polar histogram of Hes1-Achilles oscillation phase at each indicated time around division, radial axis representing count frequency.

**Extended Data Fig. 9.**
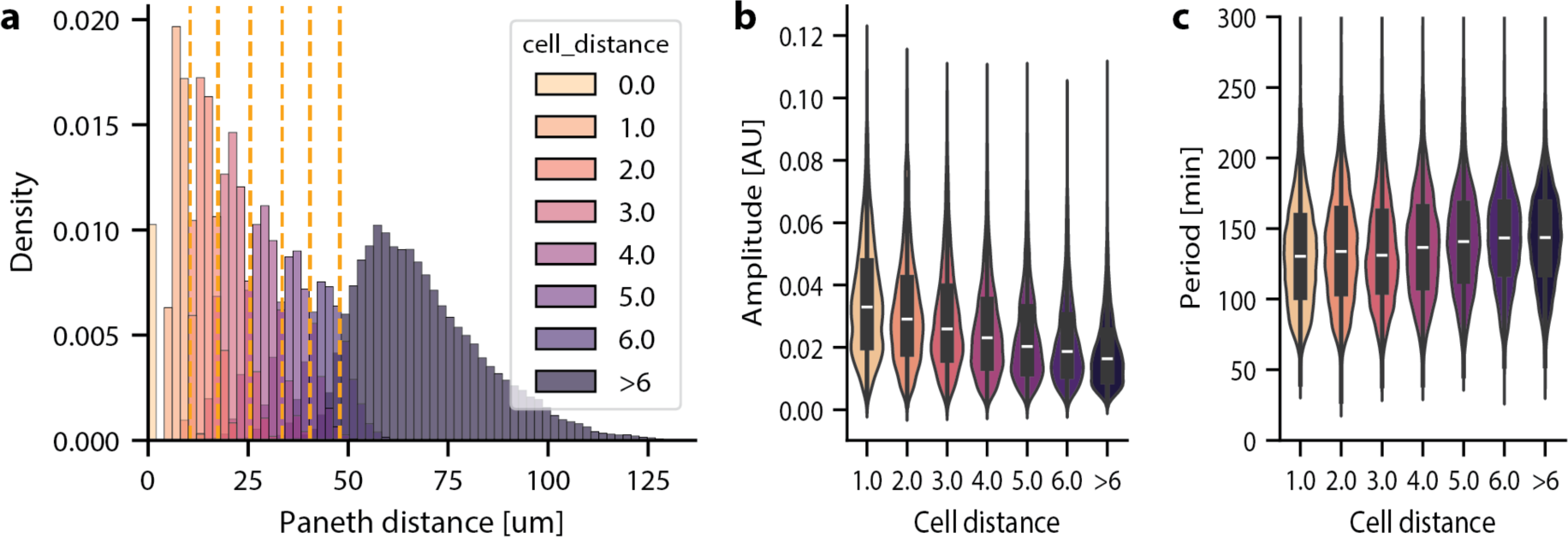
Analysis of single-cell tracks with distance from crypt centre. Intestinal organoids expressing Hes1-Achilles and H2B-mCherry were cultured in 2D, followed by live-imaging and analysis. (a) Histogram showing the distribution of cells from Paneth cells, n = 308156 cells for all timepoints. (b) Violin plot of amplitude vs distance from Paneth cells. (c) Violin plot of period vs distance from Paneth cells. n=1901 tracks, N=3 biological replicates, n_inst_ = 147959 instantaneous measurements. Normal distribution was assessed by Shapiro Wilk test, p<0.05 for all dat1asets. Significance was tested by Mann-Whitney U test, with all other cell distances compared against cell distance = 1. All comparisons were statistically significant, p<0.05.

**Extended Data Fig. 10.**
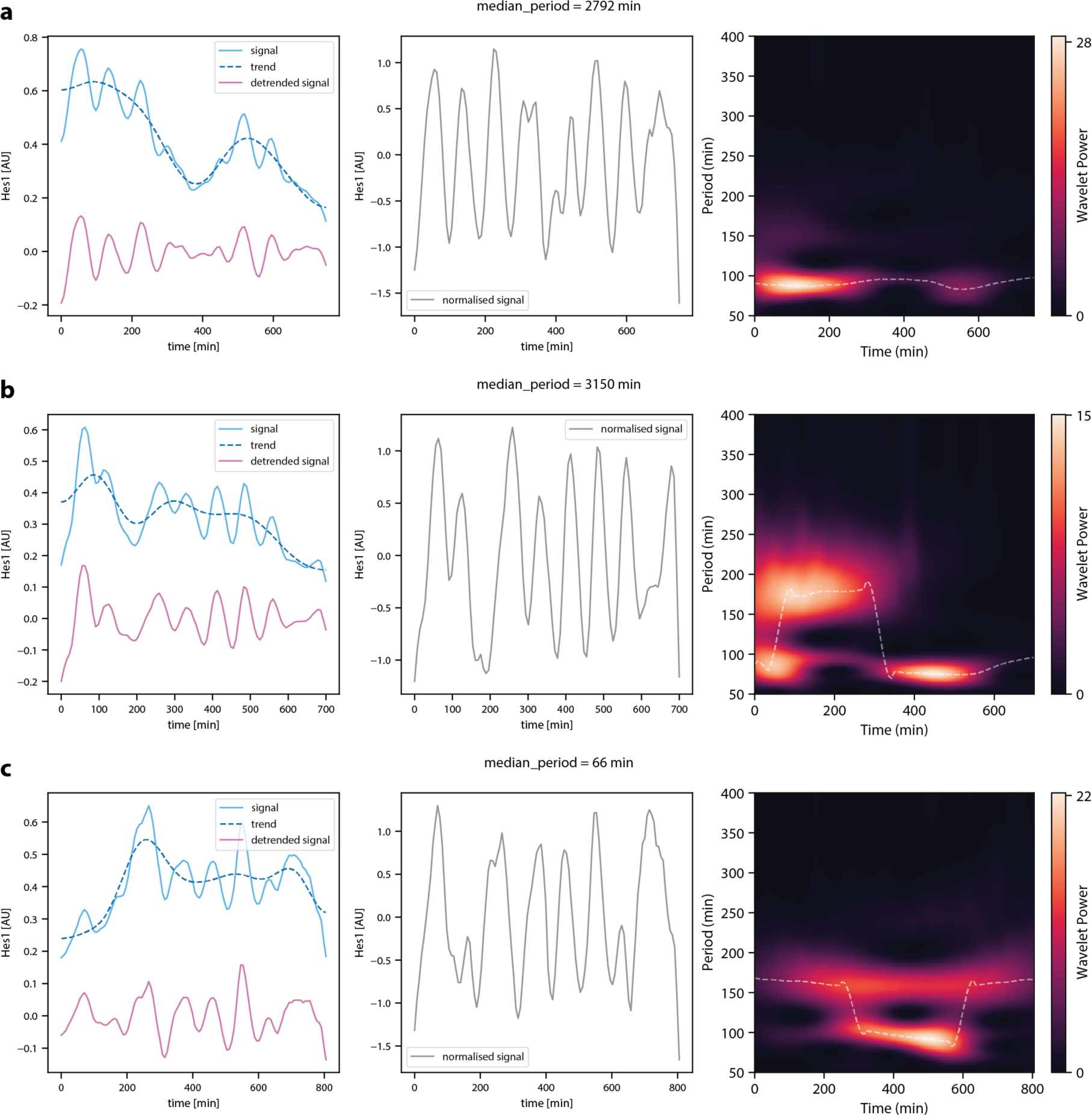
Examples of wavelet power spectra. Intestinal organoids expressing Hes1-Achilles and H2B-mCherry were cultured in 2D, followed by live-imaging and analysis. (left) Time traces of smoothed (original) Hes1-Achilles intensity signal, trend line and detrended signal. (centre) Time trace of amplitude normalised Hes1-Achilles signal. (right) Wavelet power spectra calculated from amplitude normalised Hes1-Achilles signal.

**Extended Data Fig. 11.**
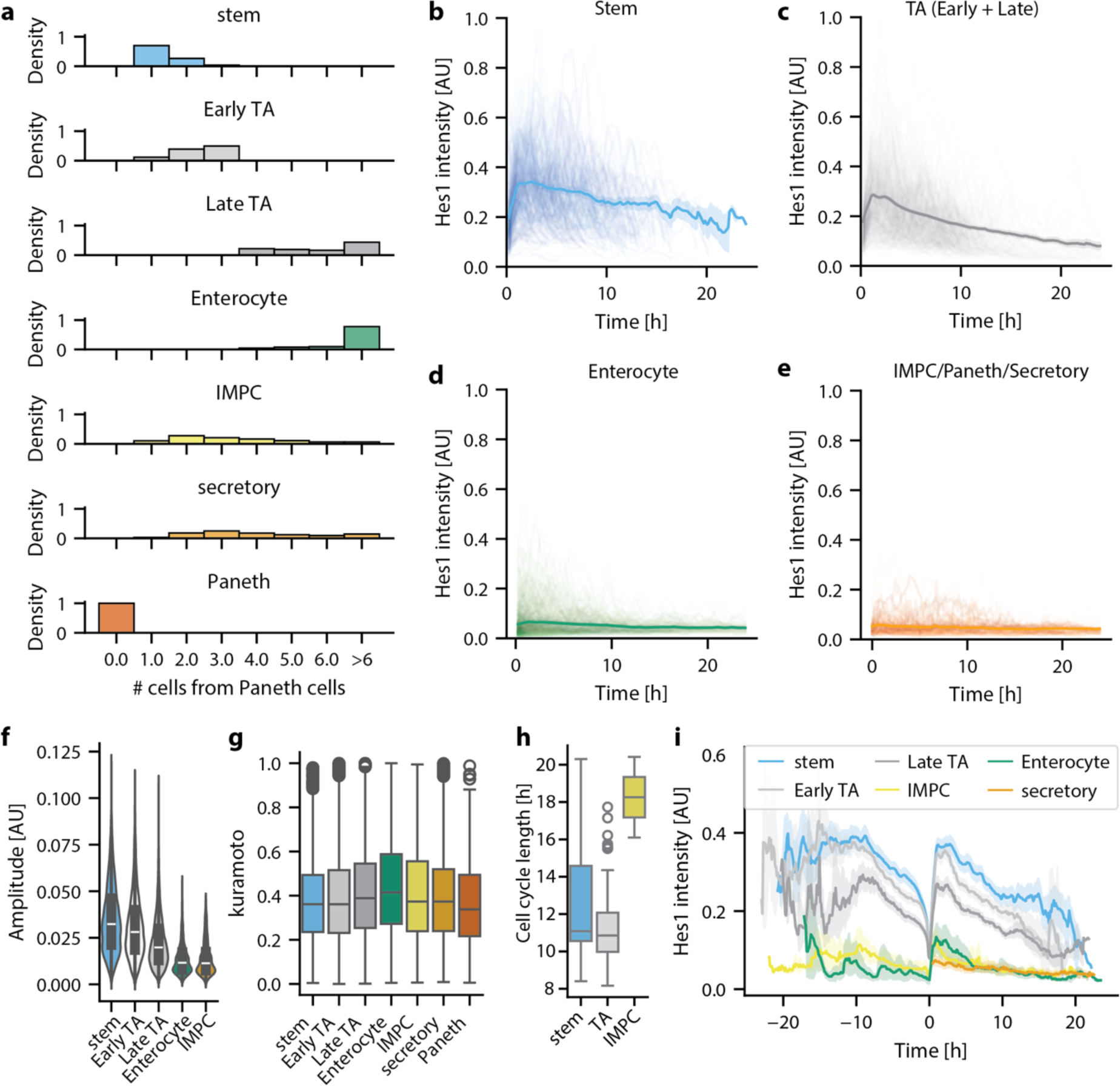
Further analysis of cell-type specific Hes1 dynamics. Intestinal organoids expressing Hes1-Achilles and H2B-mCherry were cultured in 2D, followed by live-imaging and analysis. (a) Histogram showing distribution of distances of different cell types from Paneth cells, n = 308156. (b-e) Hes1-Achilles intensity vs time for different cell types, n= 1901 tracks. (f) Violin plot of amplitude vs cell type. (g) Boxplot of Kuramoto order parameter per cell type. For (b-g), n = 1901 tracks. (h) Boxplot of cell cycle length per cell type, n = 182 tracks. (i) Hes1-Achilles intensity vs time for all dividing cells per cell type, centred around division, n = 905 tracks.

**Extended Data Fig. 12.**
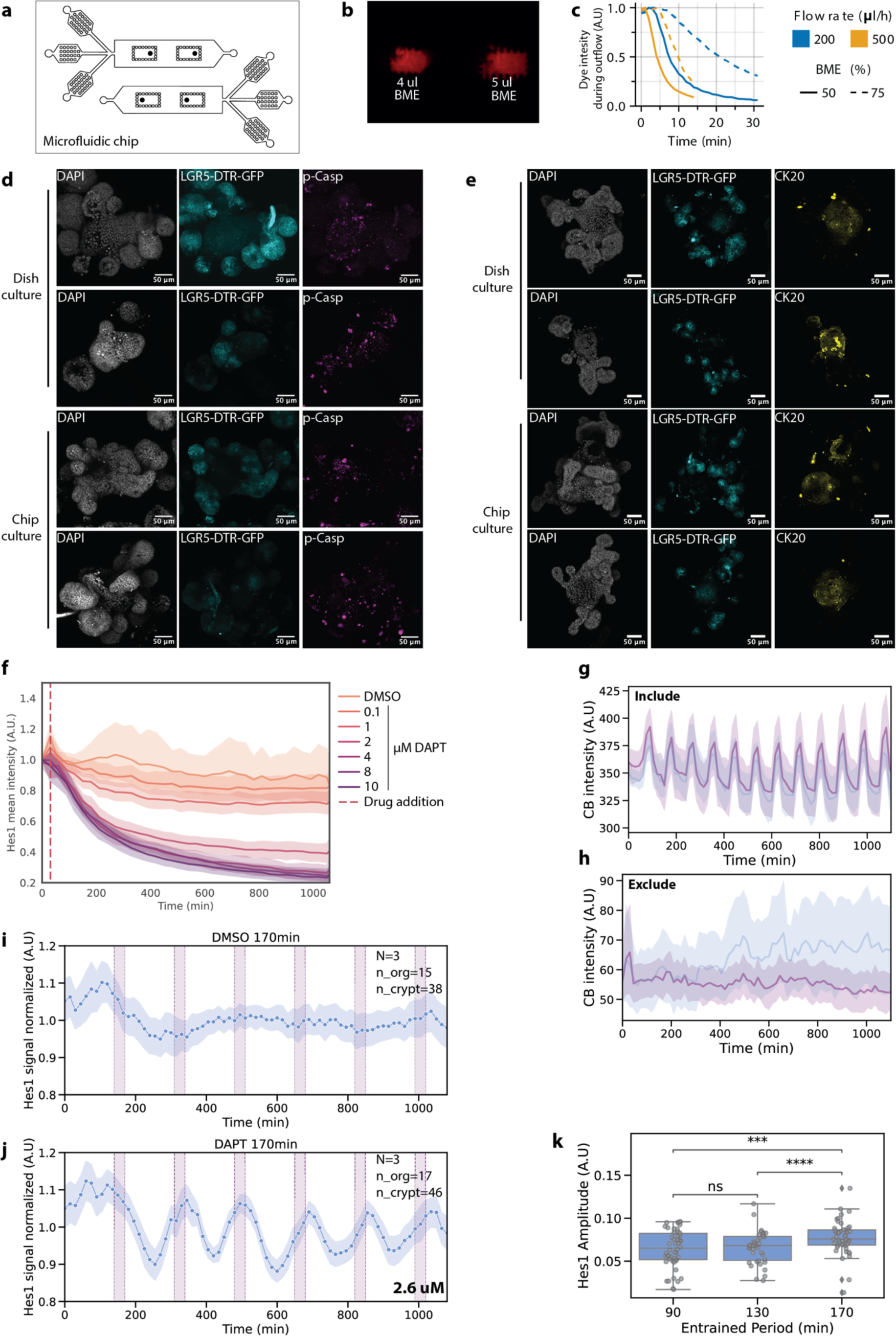
Optimization of modulating oscillations in intestinal organoids on microfluidic chips. (a) Schematic representation of chip design, two separate chambers for control (DMSO) and test (DAPT) conditions. Two organoid loading sides, three inlets and one outlet per condition. (b) Optimization of BME loading amount, fluorescent dye was added to BME. 4µl stayed within the bubble traps whereas 5µl spread through the chip (c) Dye intensity over time when following outflow of the chip. Graph indicates optimization of flow-rate and BME concentration, 200µl/h and 50% BME were chosen for the following experiments. (d) Representative images of organoids cultured on a dish versus organoids cultured on a microfluidic chip. LGR5-DRT-eGFP organoids were stained for phospho-Caspase, an apoptosis marker, and counterstained with DAPI. (e) Representative images of organoids cultured on a dish versus organoids cultured on a microfluidic chip. LGR5-DRT-eGFP organoids were stained for CK20, an enterocyte marker, and counterstained with DAPI. (f) DAPT titration from 0.1uM-10uM, on Hes1-Achilles organoids. Hes1-Achilles intensity was detected over time per crypt and normalized to the first timepoint (untreaded). Red dotted line represents treatment start time of 30 min. N=3, n_organoid= 9, n_crypt = 33 +/-3.7. (g-h) Representation of Caspase Blue (CB) intensity over time for two different microfluidic experiments. (g) Experiments that were included, (h) experiments that were excluded. (i) Entrainment of Hes1-Achilles oscillations to periodic pulses of 170-min of the NOTCH inhibitor DAPT (concentration indicated). Quantification of (normalized) Hes1-Achilles signal in organoid crypts. Experiments (N = independent experiments, n_org = individual organoids, n_crypt = individual crypts) were combined using the external force (pink bars) as an objective time reference. (j) Quantification of amplitude by wavelet transform and comparison between entrainment periods with following DAPT conditions; 90min = 1.38µM DAPT, 130min = 2µM DAPT, 170min = 2.6µM DAPT. Normal distribution was calculated by Shapiro Wilk test and significance was tested by independent t-test or Mann-Whitney U test depending on normal distribution and Bonferroni correction was performed to take multiple samples into account. ns: 5.00e-02 < p <= 1.00e+00, *: 1.00e-02 < p <= 5.00e-02, **: 1.00e-03 < p <= 1.00e-02, ***: 1.00e-04 < p <= 1.00e-03, ****: p <= 1.00e-04.

**Extended Data Fig. 13.**
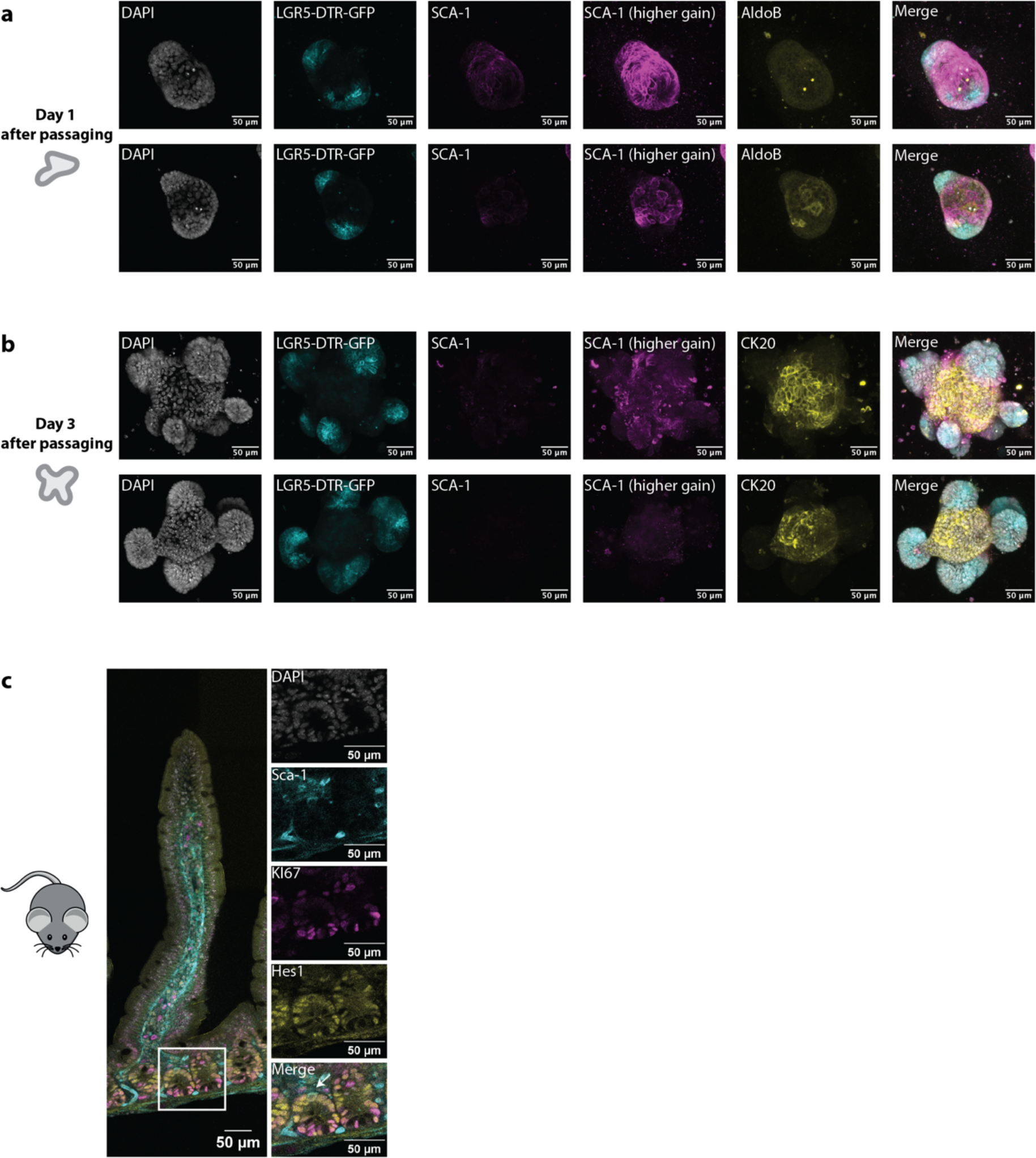
Organoid size and cell composition after passaging showing the fetal-like stem cell marker SCA. (a, b) Representative maximum intensity images of organoids 1 day (a) or 3 days (b) after passaging. LGR5-DTR-GFP organoids were stained for a fetal-like stem cell marker, SCA-1, enterocyte marker AldoB or CK20 and counterstained with DAPI. SCA-1 was depicted with a low and high threshold based on low, but present SCA-1 intensities at 3 days after passaging. (c) Representative image of Hes1-Achilles gut-slices, stained with a proliferation marker KI67, fetal-like stem cell marker, SCA-1, and counterstained with DAPI. Magnification of the crypt is shown and SCA-1 positive cell is indicated by an arrow.

**Extended Data Fig. 14.**
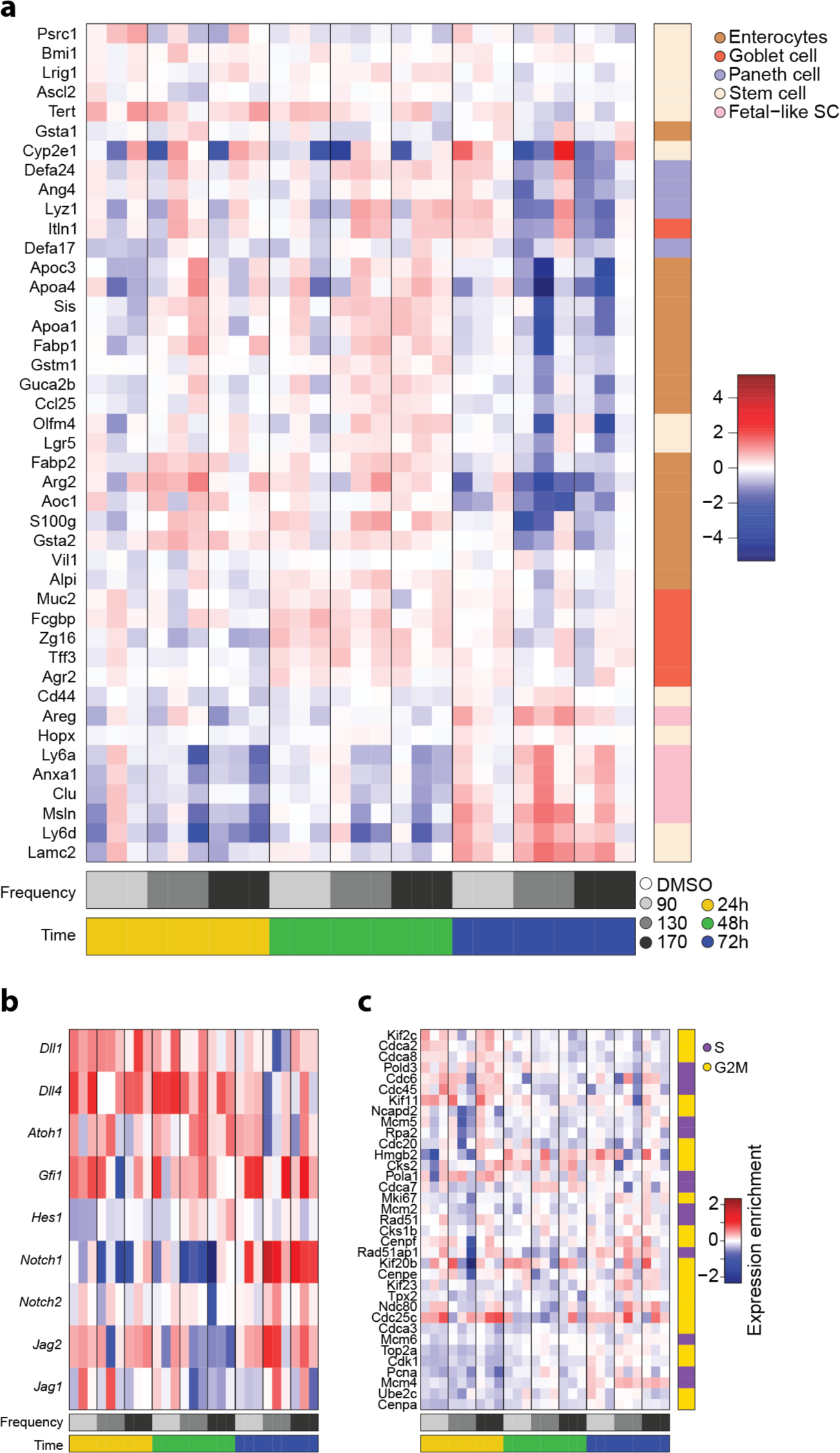
Analysis of bulk RNAseq data after entraining to different periods and durations. Hierarchical clustering of gene expression profiles of cell type specific genes (a) gene expression correlation profiles related to the NOTCH pathway (b) or to cell cycle activity (c), subdivided by entrainment and duration of entrainment. Z-score is shown. Values represent fold change relative to time-matched DMSO controls.

**Extended Data Fig. 15.**
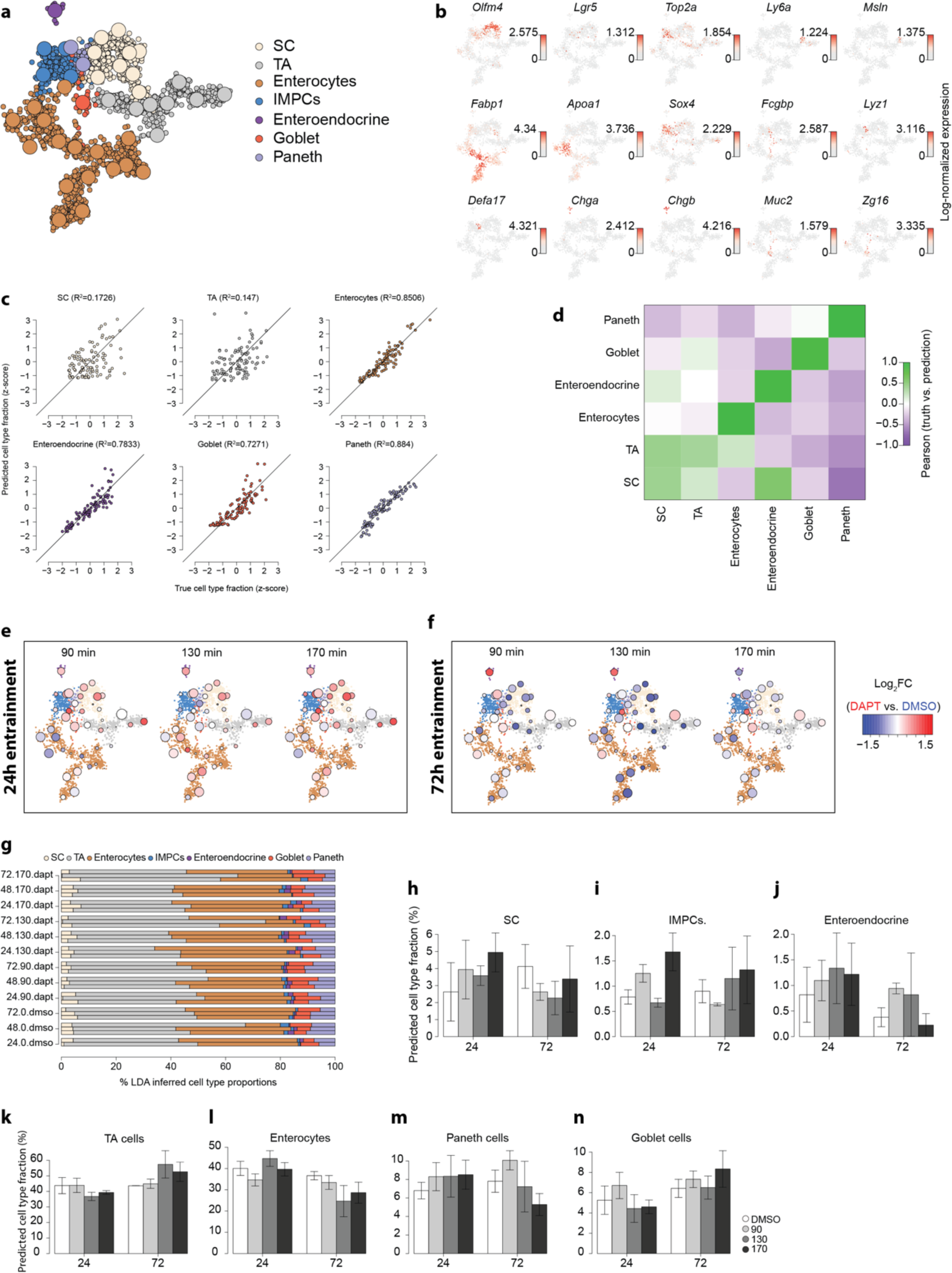
Effect of entrainment on intestinal organoids analysed by bulk RNA sequencing and Latent Dirichlet Allocation (LDA) analysis. (a) Clustering re-analysis of single-cell RNAseq data presented in Bues et al.,2022^40^. Dots represent single cells and circles represent metacells^60^. Cells and metacells are coloured by assignment to cell types. (b) Expression of lineage markers across the reanalyzed single-cell RNAseq data as in (a). Dot color indicates log normalized expression. (c, d) Validation of LDA predictions across datasets and sequencing platforms. Single-cell RNAseq data from Haber et al. (2017)^12^ was projected on the data from Bues et al. (2022)^40^. Then 100 pseudo-bulk aggregates of 200 cells each were generated and analysed via the LDA pipeline (see Methods for further details). (c) Predicted cell type proportions (z-score; *y* axis) are plotted against their true value per aggregate (z-score, *x* axis). R2 values are indicated per cell type. (d) Pearson correlation values between predicted and true cell type proportions are shown. (e, f) Results of the LDA deconvolution of the 24h (e) or 72h (f) bulk RNAseq data on top of the single-cell clustering analysis as in (a). Metacells are coloured by fold-change in their predicted proportions compared to the time-matched DMSO controls. (g) Predicted cell type distributions for each sample included in the bulk RNAseq data, based on the LDA analysis. (h-n) Predicted percentage of stem cells (h), secretory progenitors (i), enteroendocrine cells (j), TA cells (k), Enterocytes (l), Paneth cells (m) and Goblet cells (n) per entrainment vs control (DMSO) per experimental duration including 24 and 72 hours, based on the LDA analysis.

**Extended Data Fig. 16.**
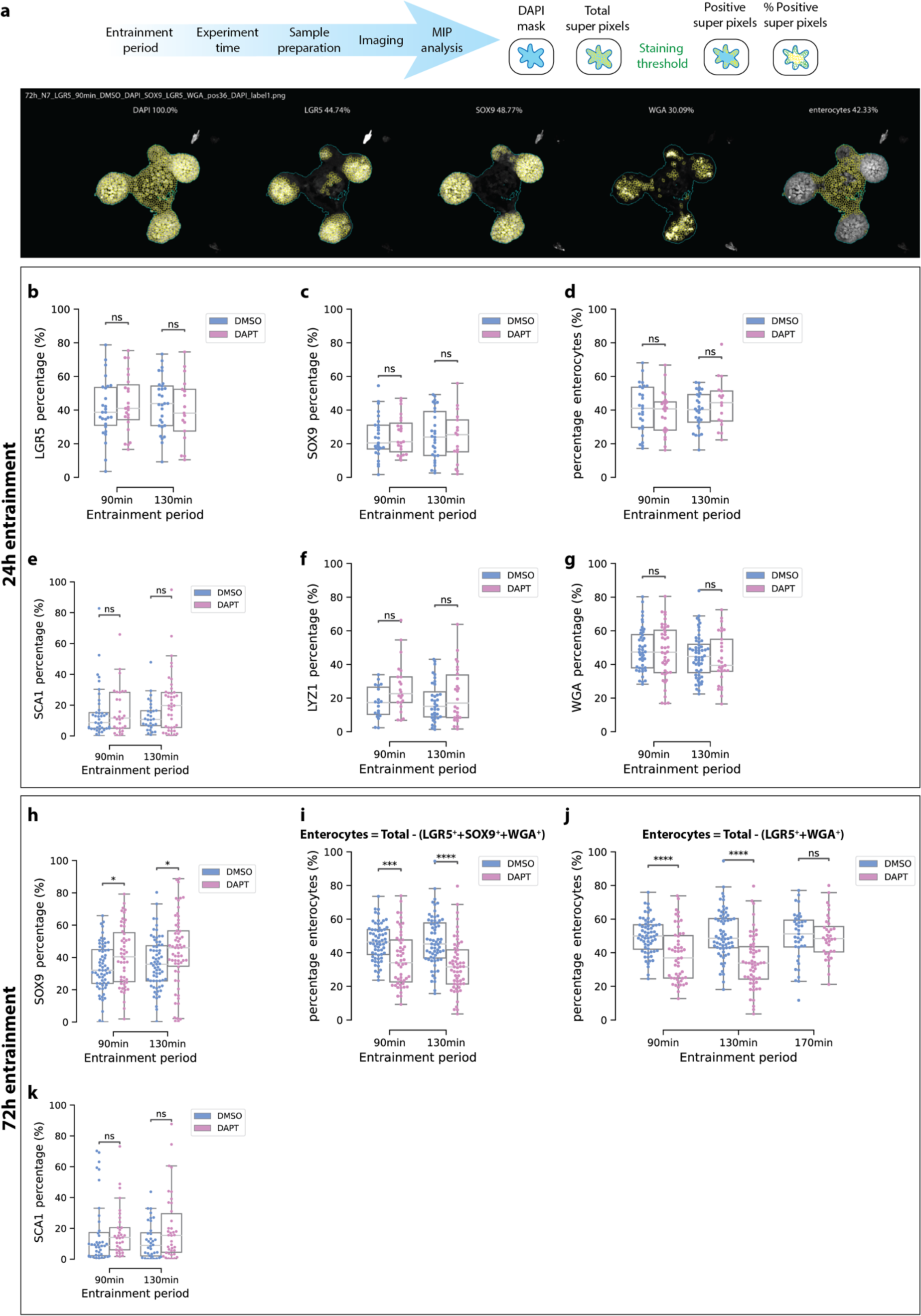
Cell type composition of intestinal organoids after entrainment. (a) *Upper panel:* Schematic representation of experiments, sample processing and image analysis. *Lower panel:* Representative images of DAPI mask and superpixel calculation. (b-g) Quantitative analysis of cell percentage per organoid compared to control (DMSO) for 90-,130-min entrainment for 24 h. Cell percentages tested for LGR5 (b), SOX9 (c), Enterocytes (d), SCA-1 (e), LYZ1 (f), WGA(g). (h-k) Quantitative analysis of cell percentage per organoid compared to control (DMSO) for 90-,130-(h, i, k) and 170-min (j) entrainment for 72 h and representative images including percentage. Cell percentages tested for SOX9 (h), Enterocytes (I, j), SCA-1 (k). Enterocytes in (j) were calculated without SOX9, so 170-min entrainment could be included. N= minimum of 3 and n_org = minimum of 14 organoids per condition. Normal distribution was calculated by Shapiro Wilk test and significance was tested by independent t-test or Mann-Whitney U test depending on normal distribution. ns: 5.00e-02 < p <= 1.00e+00, *: 1.00e-02 < p <= 5.00e-02, **: 1.00e-03 < p <= 1.00e-02, ***: 1.00e-04 < p <= 1.00e-03, ****: p <= 1.00e-04.

**Extended Data Fig. 17.**
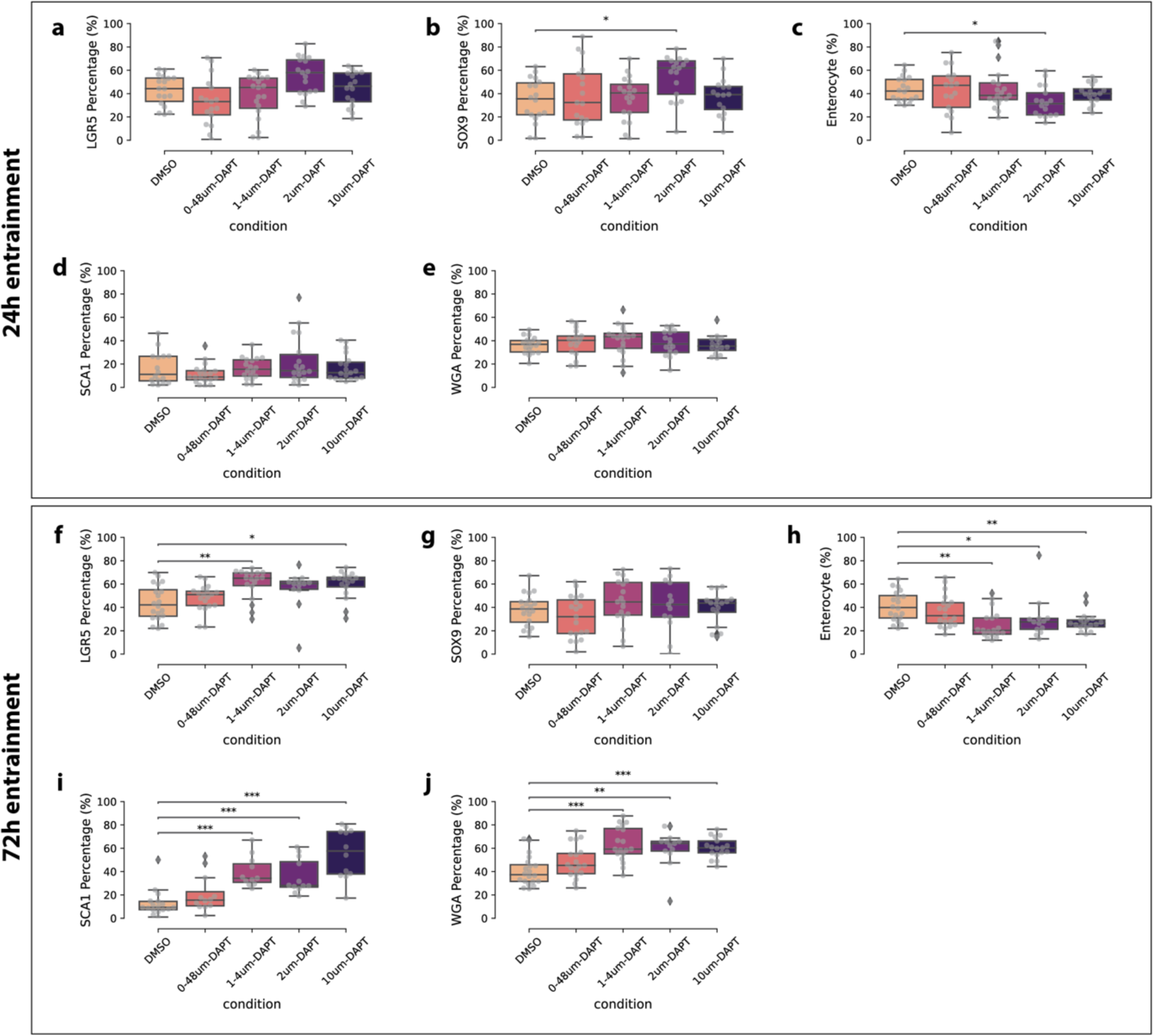
Cell type composition after constant DAPT treatment. Quantification of cell percentage per organoid by superpixel analysis when organoids are treated with 0,48µM, 1.4µM, 2µM and 10µM compared to control (DMSO) for a period of 24 hours (a-e) or 72 hours (f-j). Organoids were treated the same as the microfluidic experiments, passaged 1 day before a 72 h experiment and 3 days before a 24 h experiment. LGR5-DTR-GFP (a, f) or WT organoids were used and stained for SOX9 (b, g) a proliferation marker. Enterocytes were calculated by total superpixels – (LGR5+SOX9+WGA superpixels) (c, h). Organoids were stained for SCA-1 (d, i) a fetal-like stem cell marker or WGA (e-j) a secretory cell type marker. Normal distribution was calculated by Shapiro Wilk test and significance was tested by independent t-test or Mann-Whitney U test depending on normal distribution. Due to multiple samples Bonferroni correction was performed. ns: 5.00e-02 < p <= 1.00e+00, *: 1.00e-02 < p <= 5.00e-02, **: 1.00e-03 < p <= 1.00e-02, ***: 1.00e-04 < p <= 1.00e-03, ****: p <= 1.00e-04.

**Supplementary Table 1:**
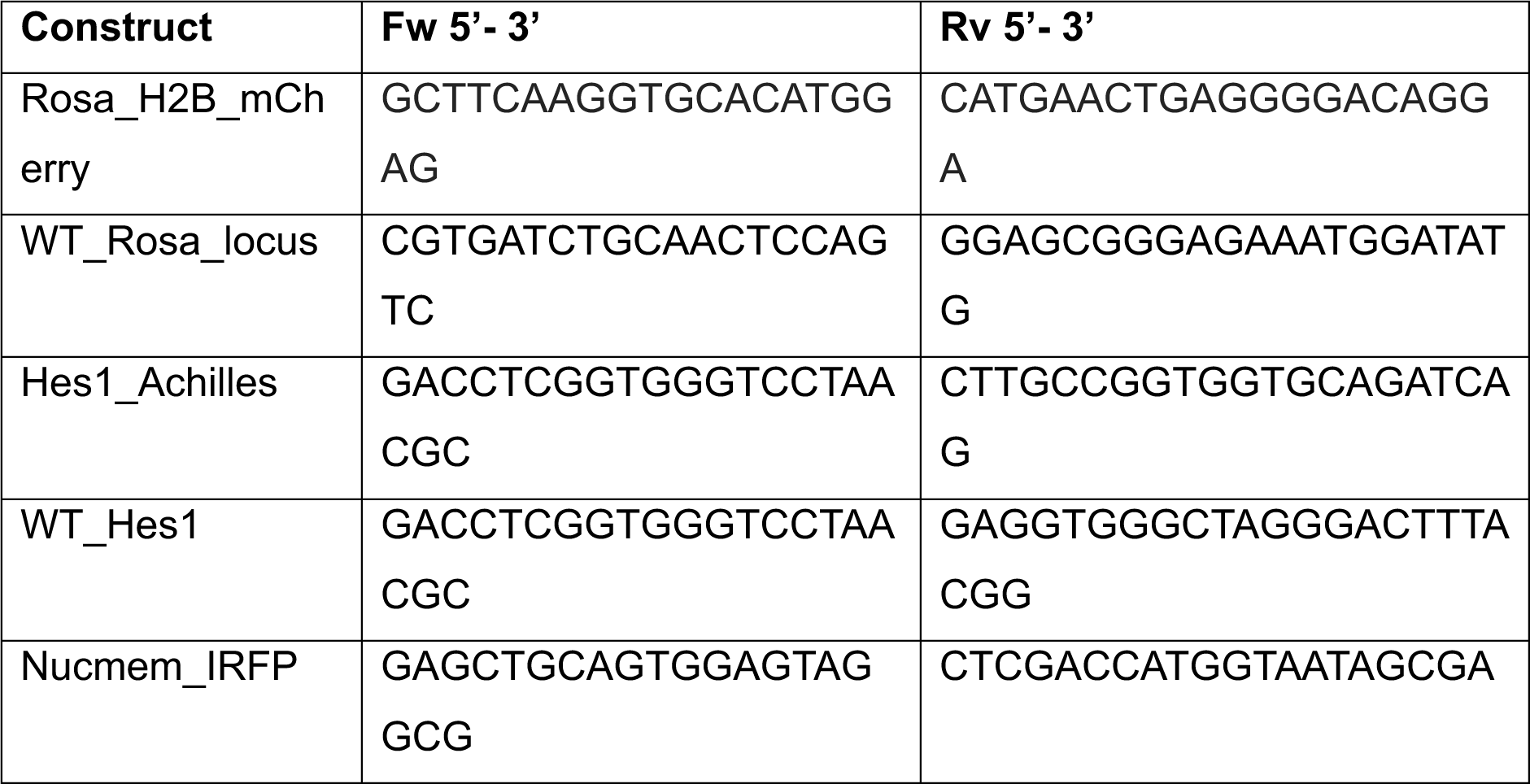
Genotype Primers.

**Supplementary Table 2:** Antibodies.

| Target | concentration | Fixation<br>time (min) | Host<br>Species | Article number |
| --- | --- | --- | --- | --- |
| Ki67 | 1:200 | 10-45 (3D) | Rat | 14-5698-82<br>(ebioscience) |
| Ki67 | 1:200 | 10-45 (3D) | Rabbit | ab16667 |
| Lyz1 | 1:200 | 45 (3D) | Rabbit | A0099 (DAKO) |
| CK20 | 1:200 | 10 (3D) | Mouse | M701929-2<br>(DAKO) |
| WGA | 1:500 | 45 (3D) | - | 29026-1<br>(Biotium) |
| Sox9 | 1:200 | 45 (3D) | Rabbit | ab185966 |
| Olmf4 | 1:200 | 20 | Rabbit | 39141 (XP) |
| Hes1 | 1:200 | 5 (3D) | Mouse | sc-166410 |
| Cleaved<br>Caspase-3 | 1:200 |  | Rabbit | 9661S(cell<br>signaling tech.) |
| AldolaseB | 1:200 | 10 (3D) | Rabbit | ab75751 |
| Muc2 | 1:200 | 20 | Rabbit | NBP1-31231 |
| ChrA | 1:200 | 45 | Mouse | sc-13090 |
| Sca-1 | 1:300 | 10 (3D) | Rat | Ab51317 |

### Supplemental movies

**Supplemental movie 1. Intravital Imaging following a single cell.** Cyan: H2B-mCherry, purple: Hes1-Achilles.

**Supplemental movie 2. Light-Sheet imaging of 3D organoid expressing Hes1-Achilles.**

**Supplemental movie 3. Imaging of 2D culture of intestinal organoids.** Purple: H2B-mCherry, cyan: Hes1-Achilles.

**Supplemental movie 4. Segmentation of 2D culture movies.** Purple: H2B-mCherry, cyan: Hes1-Achilles.

**Supplemental movie 5. Tracked single cell in 2D culture.** Purple: H2B-mCherry, cyan: Hes1-Achilles.

**Supplemental movie 6. Quantification of synchronization using Kuramoto order parameter.** *Left panel*: Cells labelled according to their Kuramoto order parameter (blue = 0, red = 1); *Middle panel*: Hes1-Achilles; *Right panel*: H2B-mCherry.

**Supplemental movie 7. Track of a dividing cell.**

**Supplemental movie 8. TA cell to enterocyte differentiation.**

**Supplemental movie 9. Stem cell to IMPC differentiation.**

**Supplemental movie 10. Imaging of organoids expressing Hes1-Achilles in microfluidic chip.** Entrainment using 130 min drug pulses. *Left panel*: DMSO control; *Right panel*: DAPT.

